# Effect of temperature on circadian clock functioning of trees in the context of global warming

**DOI:** 10.1101/2024.03.22.586279

**Authors:** Maximiliano Estravis-Barcala, Sofía Gaischuk, Marina Gonzalez-Polo, Alejandro Martínez-Meier, Rodrigo A. Gutiérrez, Marcelo Yanovsky, Nicolás Bellora, María Verónica Arana

## Abstract

Plant survival in a warmer world requires the timely adjustment of biological processes to cyclical changes in the new environment. Circadian oscillators have been proposed to contribute to thermal adaptation and plasticity in plants, due to their ability to maintain periodicity in biological rhythms over a wide temperature range, promoting fitness. However, the influence of temperature and circadian clock performance on plant behaviour in natural ecosystems is not well understood. Here we used two co-occurring *Nothofagus* tree species from the Patagonian forests that are adapted to contrasting thermal environments derived from their different altitudinal profiles. We revealed that the upper thermal limits for accurate clock function are linked to the species’ thermal niches and contribute to seedling plasticity in natural environments. We computationally identified 24 circadian clock-related genes, which showed a high degree of structural conservation with clock genes from both annual and perennial species, and very similar patterns of gene expression to those of *Arabidopsis thaliana*. Warm temperatures produced a strong transcriptomic rearrangement, which affected the expression of clock-related genes and direct clock targets, evidencing the extent of clock functioning disruption by temperature. *N. pumilio*, the species from colder environments, showed reduced ability to keep rhythmicity at high temperatures compared to *N. obliqua*, which inhabits warmer zones. Accordingly, *N. pumilio*, but not *N. obliqua*, showed a limited oscillator function in warmer zones of the forest, reduced survival, and growth. Together, our results highlight the potential role of a resonating oscillator in ecological adaptation to a warming environment.

## Introduction

Increasing worldwide temperatures are expected to have major impacts on primary forest components such as trees, and result in notable changes in the ranges (1), composition (2) and phenology (3) of species, even to the extent of creating uncertainty in climate change predictions (3, 4). A key question therefore is to understand how increased temperatures will affect pathways that regulate the expression of traits of ecological relevance in forest trees. In this regard, the effects of temperature on circadian oscillator function deserves investigation, given their pervasive role in controlling a myriad of biological processes.

Circadian oscillators, also known as circadian clocks, are entrained time-keepers that include gene networks of interlocked feedback loops. They generate rhythmic outputs (circadian rhythms) with a period usually near 24 hours, adjusting biological processes to cyclical changes in the environment caused by the earth’s rotational and orbital movements (5). They therefore function as integrators between organisms and their environments, promoting fitness (6–8) and occurring in Nature from Archaea to crown eukaryotes (9, 10). In plants, most of our knowledge of the circadian clock has been learned through forward and reverse genetics in the model plant *Arabidopsis thaliana* (11). In this annual species, the clock controls diverse functional traits throughout the life cycle, as well as cellular processes related to stress responses and gene expression (8, 11–15). By contrast, studies in perennial species such as trees are relatively scarce, and clock function has been evaluated by analysing temporal patterns of expression of core and circadian-controlled genes (16–19). In trees, the clock modulates photoperiodic control of growth (20) and canopy gas exchange to an extent that affects whole-tree water use, potentially influencing biosphere-atmosphere interactions (21, 22).

A remarkable characteristic of circadian oscillators is that they are temperature compensated, such that the period of their oscillation is less sensitive to temperatures in the physiological range than would be expected from physical laws (5). In plants, it has been proposed that this range of temperatures is species-specific, and that this characteristic may contribute to thermal adaptation and plasticity, potentially influencing responses to climate warming (22). However, the impact of temperature on circadian clock performance in natural environments, and the contribution this makes to plant growth, are not well understood. In part this is because early experiments were carried out under controlled conditions, testing the effects of constant temperatures using circadian protocols, i.e. seedlings exposed to non-variable environments after a period of training by light-dark or temperature cycles (5, 11). While such approaches provide valuable information on the effect of temperature as a single factor on the oscillator itself, they are not representative of most natural environments, where for instance, day and night constitute a permanent training signal that can potentially override disruption due to high temperature.

Here we used a multidisciplinary approach that involves experiments under both controlled and natural environments to assess whether the ability of two closely-related tree species to inhabit different thermal environments and respond to environmental temperatures can be interpreted with reference to their clocks. We addressed this question using *Nothofagus pumilio and Nothofagus obliqua* (Fig. 1), two emblematic tree species from the Patagonian forests of South America that are adapted to specific thermal environments derived from their different altitudinal profiles. They belong to Nothofagaceae, a monotypic family in the order Fagales that dominates almost all of the forestry landmass of the southernmost woody ecosystem of the world: the sub-Antarctic temperate forests of Patagonia (23). In the lab, we tested the effect of temperature as a single variable on the performance of their circadian clocks. In the forest, we addressed how the natural thermal environment, both inside and outside of their natural altitudinal distribution, related to their ability to maintain internal rhythms with daily changes in the environment, and how this was associated with plant performance. Specifically, we hypothesize that the upper thermic range for accurate clock function relates to the thermic characteristics of a species’ favoured ecological niche, and to the ability of seedlings to maintain rhythms and grow in warm environments.

**Fig. 1.**
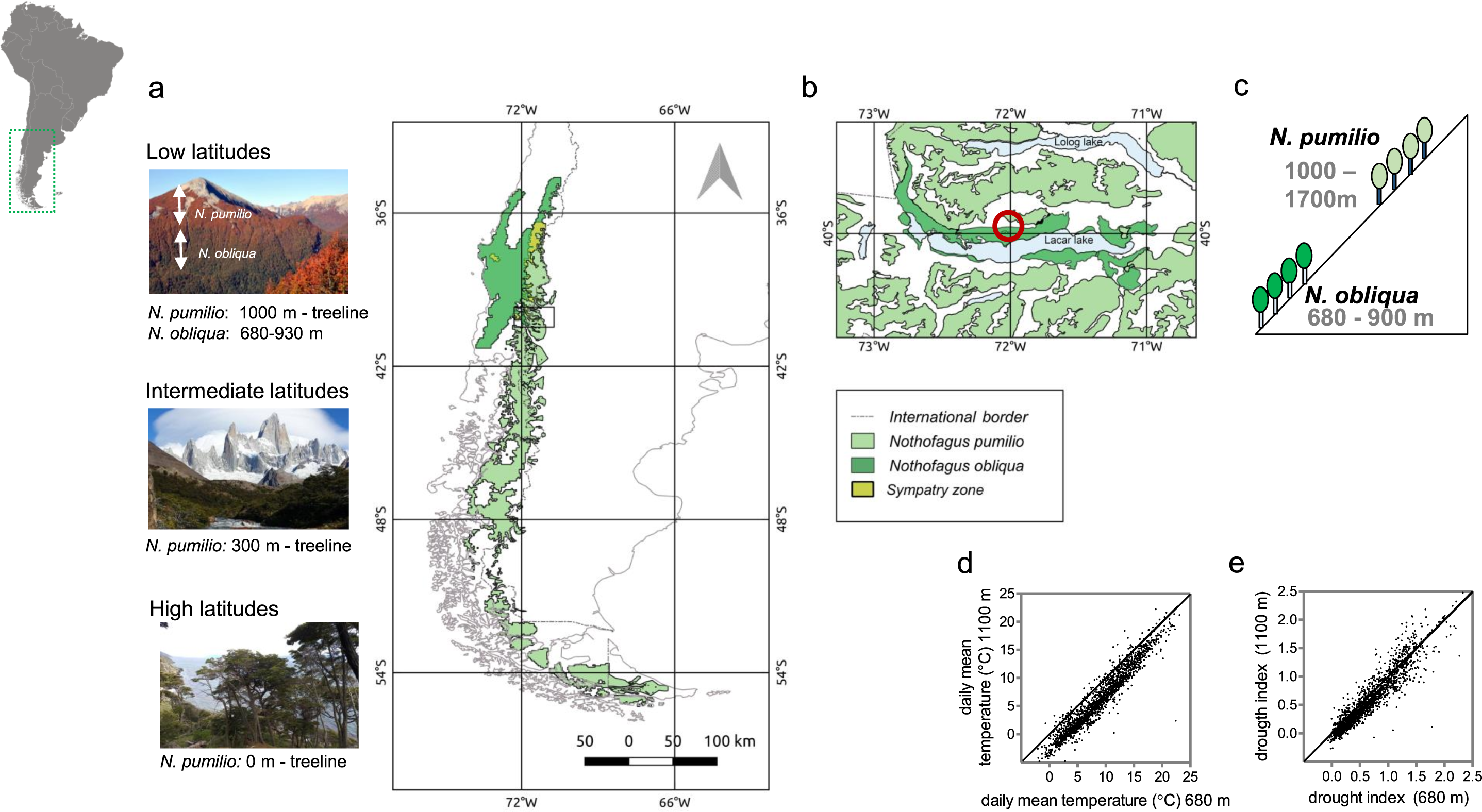
Natural distribution of the species and study site. (a) Map showing the distribution of both species. Pictures at the left show representative images of *Nothofagus pumilio* and *N. obliqua* forests, when present, at high, intermediate and low latitudes in Patagonia. The black rectangle indicates the zone that comprises the study site. (b) Study site in the Lacar lake basin, Lanin National Park, Argentina. (c) Schematic representation of the altitudinal distribution of *N. pumilio* and *N. obliqua* in the area of study. (d-e) Comparison of daily mean temperatures (d) and daily mean drought index (e) between the altitudinal sites of greatest abundance of *N. obliqua* (680 m.a.s.l.) and *N. pumilio* (1340 m.a.s.l.) during the period 2011-2017. The drought index was calculated as ((3 * Tmax) + Tmin)/(1 + HRm), where Tmax is the daily maximum temperature, Tmin is the daily minimum temperature and HRm is the daily average air humidity. Upper left: Map of South America. The green rectangle indicates the zone shown in (a).

## Results

### Response of the circadian oscillator to warm temperatures under constant conditions

Circadian rhythms in plants are observed at cellular and whole-organism levels (13, 24). The validity of studying circadian rhythms and expression of genes of the core oscillator as a common reference for circadian clock function is long established. This consistency is largely due to the fact that these phenotypes show daily rhythmicity that takes the form of sinusoidal waves with specific phases, and changes or loss of phase mirror changes in central oscillator function (25). We first investigated the impact of temperature on the functioning of the core oscillator of *Nothophagus pumilio*, chosen because it is the native tree species with the widest distribution in the Patagonian forests. It ranges in elevation from 0 to 2000 meters above sea level (m a.s.l) in high latitudinal zones (55°S), but occurs only in the coldest, sub-alpine zone above 1000 m a.s.l in warmer regions north of 41°S (23). This distribution suggests that it lacks adaptation to warmer environments and highlights a potential susceptibility to global warming. Genomic information regarding clock components is not available for this or comparable tree species, so we computationally identified the genes of the circadian clock from a draft assembly of its nuclear genome. We identified 24 circadian clock-related genes, which showed a high degree of structural conservation with related genes in *A. thaliana*, *Populus trichocarpa* and *Betula pendula* (Fig. 2, Figs. S1-2, Table S10). This information allowed us to unambiguously choose loci representing core genes to assay clock performance at the molecular level (Fig. S3). We chose 7 genes of the core oscillator that (1) were previously described in other plant species as central to core clock function, and (2) showed temporal windows of maximum expression that were distinguishable from one another, including morning, noon, evening and night. We selected the orthologues to *LATE HYPOCOTYL ELONGATION* (*NpLHY*), with a maximum of expression reported in the morning; *PSEUDORESPONSE REGULATOR 5* (*NpPRR5*) and *GIGANTEA* (*NpGI)*, with a maximum reported in the afternoon; *PSEUDORESPONSE REGULATOR 1* (*NpTOC1*), *EARLY FLOWERING 3* (*NpELF3*) and *LUX* (*NpLUX*) which peak during the evening; and *REVEILLE 1* (*NpRVE1*) with maximum expression late in the night and in the early morning (5, 11). These genes, with the exception of *NpGI* and *NpELF3* that didn’t belong to multi-genic families, clustered together with their described orthologues in model and non-model species (Fig. S4a-b), and showed a rhythmic pattern of expression in *N. pumilio* seedlings at 20°C under circadian conditions (Fig. 3, Fig. S5a-g). Peaks of maximum expression coincided with those described in *A. thaliana* (Fig. S6). The observed oscillation patterns were severely affected at 34°C (Fig. 3, Fig. S5h-n), indicating that functioning of the circadian clock of *N. pumilio* is disrupted by warm temperatures.

**Fig. 2.**
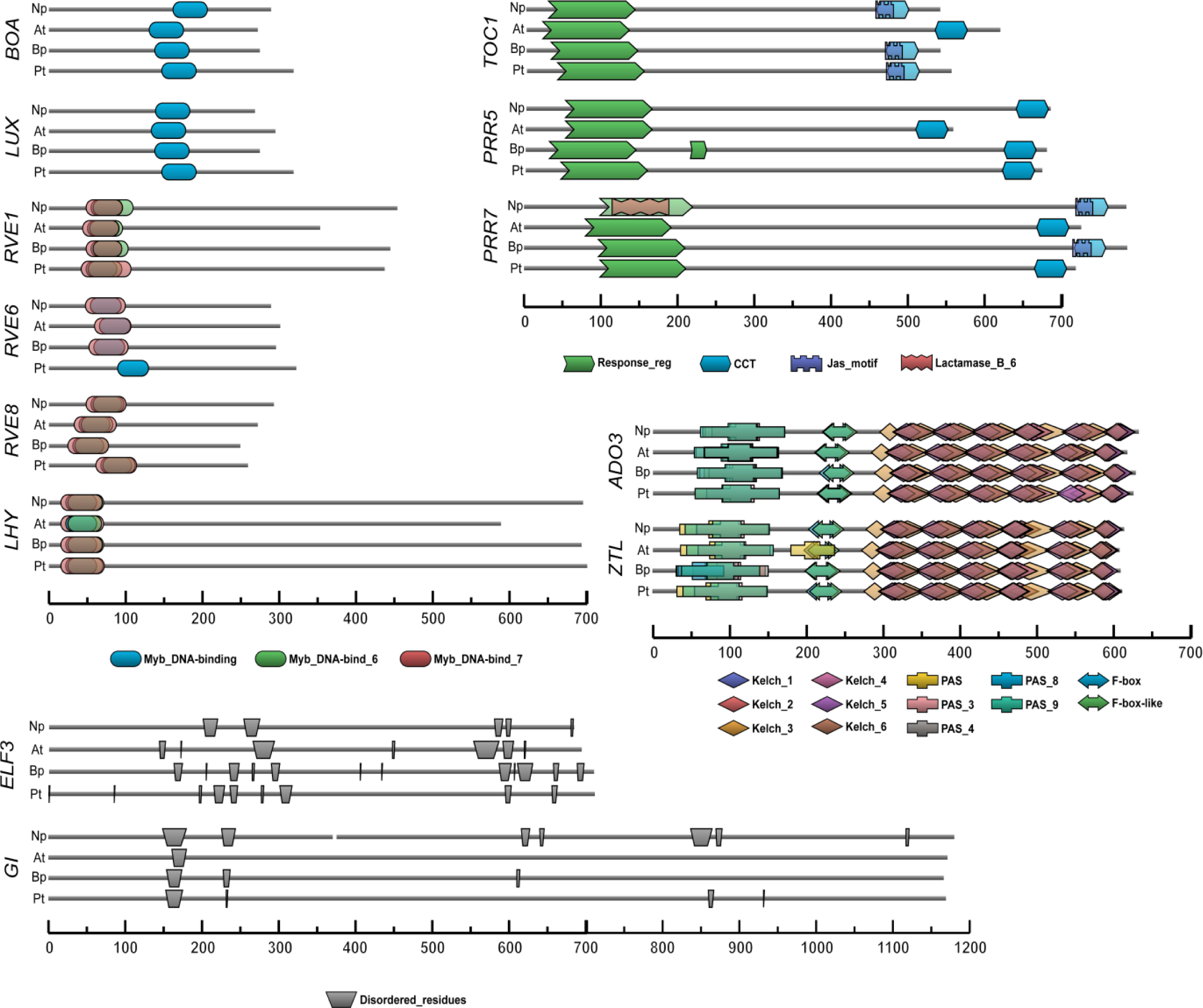
In silico characterization of core circadian clock genes. Proteins with a primary function in the circadian clock mechanism are shown. Pfam protein domains were predicted using HMMER. Positions and *p*-values of domains are listed in Table S10. In proteins lacking predicted Pfam domains (GI and ELF3), disordered regions were predicted using flDPnn (Fig. S2). The scale bar represents the amino acid number. Np: *Nothofagus pumilio*. At: *Arabidopsis thaliana*. Bp: *Betula pendula*. Pt: *Populus trichocarpa*.

**Fig. 3.**
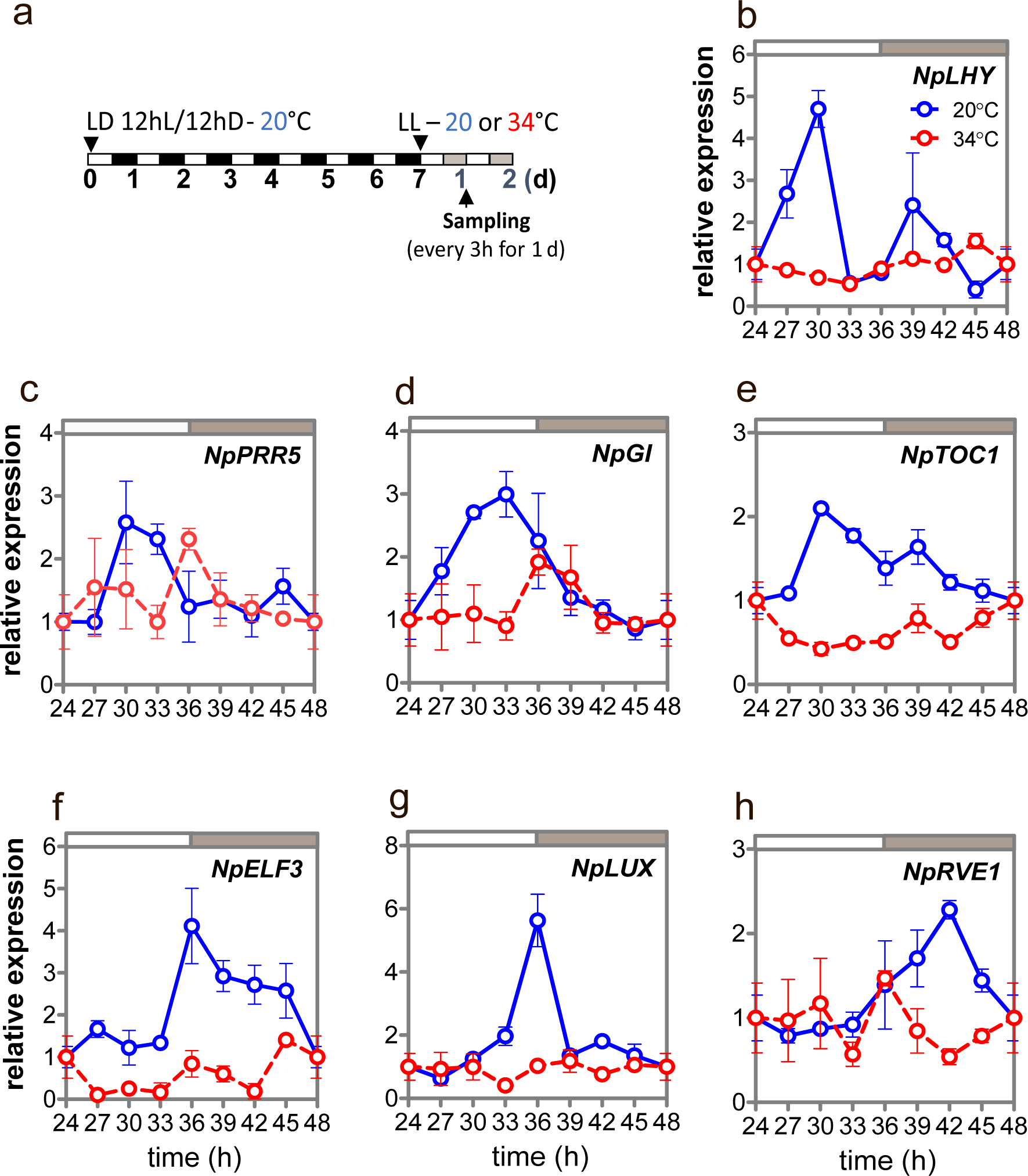
Influence of temperature on the expression of clock genes under circadian conditions. (a) Experimental protocol. Seedlings were entrained with 12h light / 12h darkness photocycles at 20°C and then exposed to continuous light (free running conditions) at 20°C or 34°C. Samples were taken every 3h during the second day after release into continuous light. (b-h) Expression of *NpLHY*, *NpPRR5*, *NpGI*, *NpTOC1*, *NpELF3*, *NpLUX* and *NpRVE1* at 20°C (blue symbols) or 34°C (red symbols). Data represent mean and SD of three technical replicates. Experiments were repeated twice with similar results (Fig. S5); results from one representative experiment are shown. Gene expression levels at ZT0 were set to 1 and used as reference for all other time points. White and grey rectangles represent subjective day and night, respectively.

To better understand the extent of warm temperature disruption of clock functioning, we studied the global patterns of gene expression of *N. pumilio* seedlings exposed to circadian conditions at 34°C. We used plants harvested 48h and 60h after release into continuous light, encompassing the periods of subjective dawn and dusk respectively (i.e., when lights would turn on or off during entrainment) (Fig. 4a). As a control we used the transcriptome of seedlings exposed to 20°C, where the circadian oscillator functions normally (Fig. 3). We found that differentially expressed genes between subjective dusk and dawn at 20°C (Fig. 4b) were enriched in circadian-regulated orthologues of *Arabidopsis thaliana* (Fig. 4c-d). Moreover, genes upregulated at subjective dawn and dusk showed enrichment in *Arabidopsis* orthologues up-regulated in early phases of day and evening, respectively (Fig. 4e-f). Treatment with 34°C reduced the amount of differentially expressed genes between subjective dawn and dusk (Fig. 4b, Tables S2-3). Warm temperatures severely affected the time-of-day patterns of gene regulation, given that 72.3% of the differentially expressed genes between subjective dawn and dusk at 20°C lost their regulation at 34°C (Table S4). Genes differentially expressed only at 20°C (72.3%) or at both 20°C and 34°C (28.5%) were enriched in circadian-regulated *Arabidopsis* orthologues (Fig. 4g-h) and in Gene Ontology biological processes (GO-BP) such as cell wall biogenesis and growth (Tables S5-6). By contrast, genes that were differentially regulated exclusively at 34°C (56% of all the genes regulated at 34°C) did not show an overrepresentation of circadian-regulated orthologues (Fig. 4i), and were enriched in GO-BP related to stress response, defence and diterpenoid biosynthesis among others, suggesting they are regulated by heat stress (Table S7). To study the effect of disruption of clock functioning by warm temperatures exclusively on orthologues of direct clock targets, we searched for *Arabidopsis* orthologues that (1) are direct TOC1 targets (26), and (2) were differentially expressed between times of the day at 20°C. Then we analysed their expression at 34°C (Fig. 4j, Table S8). We found that 34°C caused mis-regulation or a decrease in the differences of time-of-day expression of 72.5% of those genes differentially expressed at 20°C, indicating that warm temperatures provoke a general disruption of the regulation of the expression of clock targets. Moreover, genes that showed an increase in the amplitude of expression between times of the day at 34°C (22.5%) were enriched in the GO-BP ‘detoxification’ (Table S9), suggesting that they were mostly regulated by thermal stress.

**Fig. 4.**
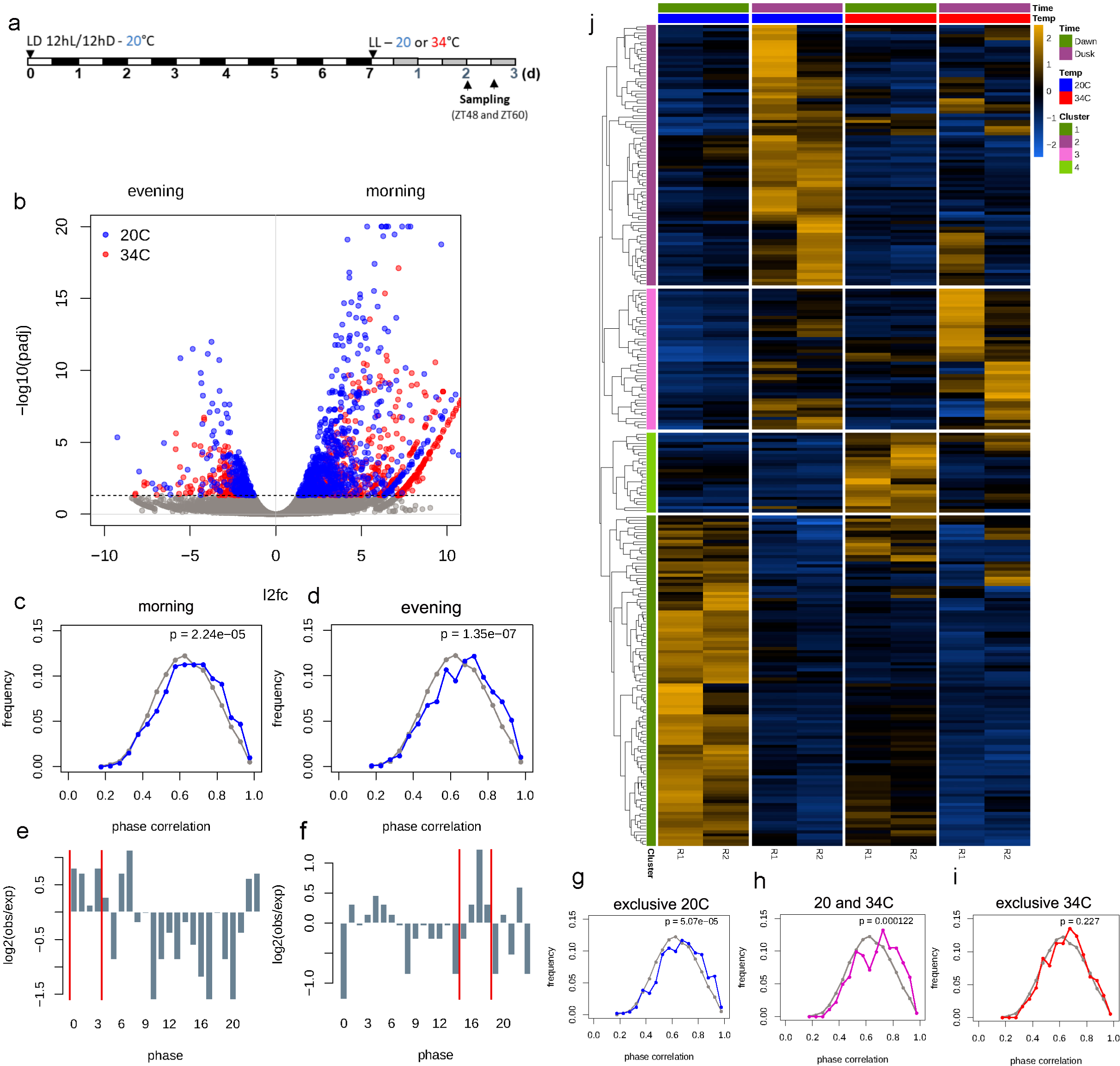
Genome-wide regulation of gene expression in circadian conditions. (a) Experimental protocol. Seedlings were entrained with 12h light / 12h darkness photocycles at 20°C and then exposed to continuous light at 20°C or 34°C. Samples were taken 48h and 60h after release into continuous light at either 20°C or 34°C. ZT refers to the time elapsed under free-running (constant light) conditions. (b) Volcano plot showing differentially expressed genes between subjective dawn (left) and dusk (right) at 20°C or 34°C. Significant expressed genes (*p* < 0.05) are shown with blue (20°C) or red (34°C) circles. (c-d) Comparison between the expected (grey) and observed (blue) frequency distribution of genes with MBPMA values between 0 and 1. We used the MBPMA cut off of 0.8 as the threshold of cycling. (c) genes over-expressed at subjective dawn, (d) genes over-expressed at subjective dusk. (e-f) log2 fold change of observed vs. expected ratio of the number of genes arranged by their time-of-day expression (phase). Phases of expression were assigned using the information of *Arabidopsis thaliana* putative orthologues that were strongly cycling (MBPMA cut-off of 0.8) of the database LL12_LDHH available in http://diurnal.mocklerlab.org/ (see methods). (e) and (f) corresponds to *N. pumilio* orthologue genes over-expressed in the subjective dawn and dusk, respectively. (g-i) Comparison between the expected (grey) and observed frequency distributions of genes with MBPMA values between 0 and 1. We used the MBPMA cut off of 0.8 as the threshold of cycling. (g) genes exclusively regulated at 20°C (blue), (h) genes regulated both at 20 and 34°C (purple) (i) genes exclusively regulated at 34°C (red) (j) Heat map showing the expression of TOC1 direct target orthologues in *N. pumilio*.

### Response of the circadian oscillator to warm temperatures under day-night photocycles and its relation to thermal characteristics of species’ preferred ecological niches

The circadian oscillator is entrained by day and night cycles, allowing trees to perceive seasonal changes and adjust physiological processes in a timely manner (20). The training signal provided by light and darkness can therefore be a cue that resets the oscillator daily, overriding the disruption of circadian clock functioning due to high temperature. To investigate this, we analysed the effect of temperature on the functioning of the clock in *N. pumilio* seedlings exposed to photocycles (Fig. 5a). All the tested genes showed maximum peaks of expression at 20°C (Fig. 5c-i, Fig. S7a-g) with similar phases to those described for *A. thaliana* (Fig. S8). Nevertheless, the warming-induced disruption of cyclic patterns of core gene expression persisted even in diurnal conditions (Fig. 5c-i, Fig. S7h-u). We then analysed the functioning of the circadian clock in the *N. pumilio* closely related species *Nothofagus obliqua*. This species co-exists with *N. pumilio* in north Patagonia, but occupies a non-overlapping thermal niche, constituting the dominant species in warmer and lower environments c. 650 m a.s.l (23) (Fig. 1). As expected, *N. obliqua* showed a marked rhythmicity in the expression of clock-related genes at 20°C (Fig. 5k-q, Fig. S9a-g) with comparable phases to *N. pumilio* and *A. thaliana* (Fig. 5c-i, Fig. S7-8). However, contrary to *N. pumilio*, rhythmicity in gene expression was robustly maintained at 31°C (Fig. 5k-q, S9h-n), showing that the oscillator of *N. obliqua* keeps rhythms at warmer temperatures than for *N. pumilio*. At 34°C, the oscillation of at least four of the seven core genes tested was disrupted (Fig. 5k-q, Fig. S9o-u). Therefore, high temperatures, in the range of 34°C, were detrimental for the functioning of the circadian clocks of both species.

**Fig. 5.**
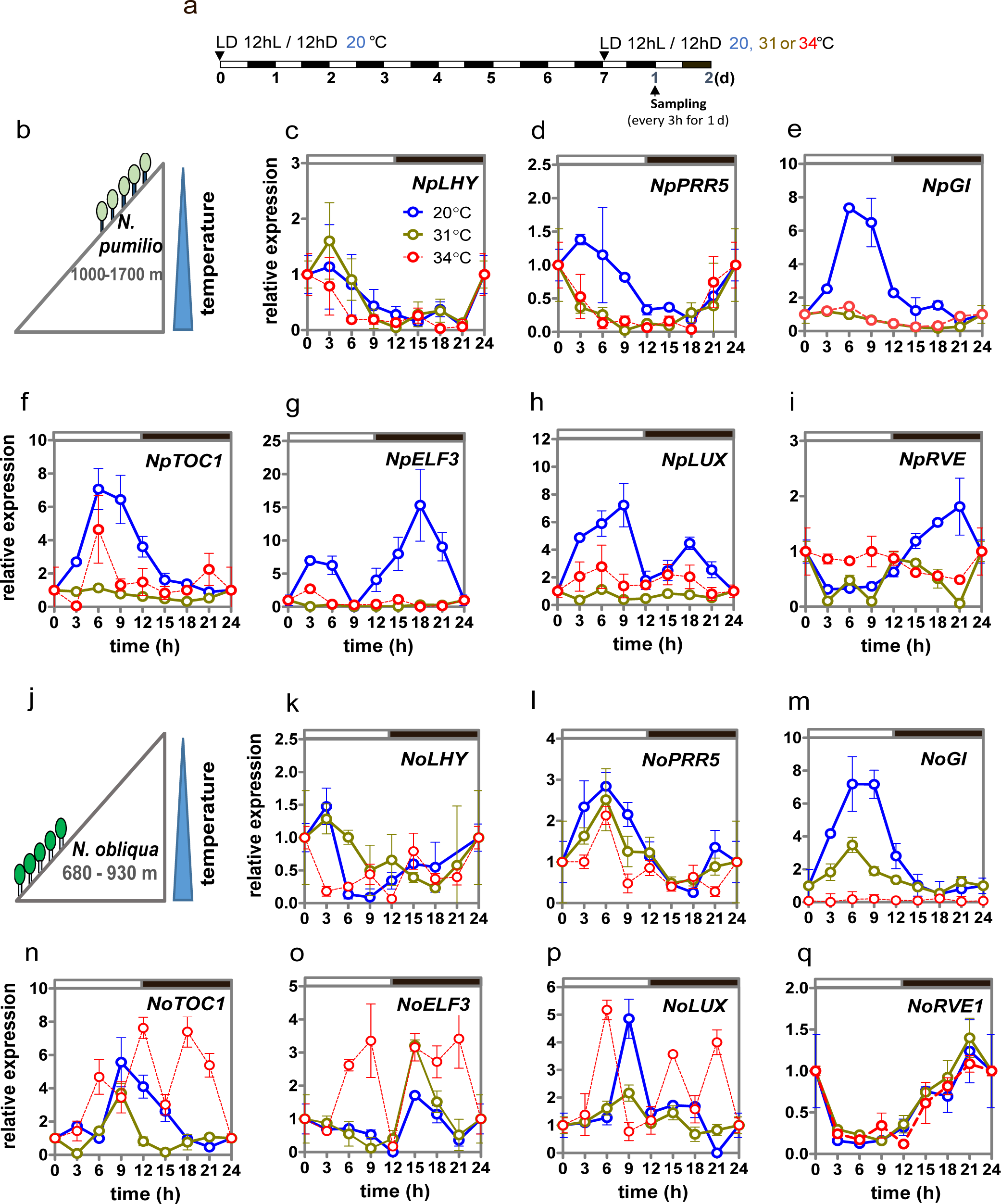
Influence of the temperature on the expression of clock genes under diurnal conditions. (a) Experimental protocol. Seedlings were entrained with 12h light / 12h darkness photocycles at 20°C and then exposed to 20°C (blue), 31°C (brown) or 34°C (red). Samples were taken every 3h during the second day of each temperature treatment. (b) and (j) schematic representation of natural distribution of *N. pumilio* and *N. obliqua*, respectively. (c-i) Expression of *N. pumilio* clock genes *NpLHY*, *NpPRR5*, *NpGI*, *NpTOC1*, *NpELF3*, *NpLUX* and *NpRVE1* at 20, 31 or 34°C.(k-q) Expression of *N. obliqua* clock genes *NoLHY*, *NoPRR5*, *NoTOC*, *NoELF3*, *NoLUX* and *NoRVE1* at 20, 31 or 34°C. Data represent mean and SD of three technical replicates. Experiments were repeated twice with similar results (Figs. S7-9); results from one representative experiment are shown. Gene expression levels at ZT0 were set to 1 and used as reference for all other time points. White and black rectangles represent day and night, respectively.

### Circadian oscillator function in altitude-swap experiments and its association to physiological traits

We aimed to investigate the extent to which observed inter-specific differences in the upper thermal ranges of accurate clock function were linked to the ability of species to grow and survive in different natural thermal niches. Using the mountain as a natural laboratory, we did altitude-swap experiments where clock and plant attributes were evaluated in seedlings in different natural thermal environments (Fig. 6a). Seedlings were transplanted into a common soil substrate to avoid any bias due to soil differences between altitudes, and were exposed to natural conditions for one month before sampling. We found that, regardless of altitude, both species showed rhythmic patterns of expression in their core genes (Fig. 6c-f, Fig. S10). Nevertheless, *N. pumilio* showed clear phase shifts between altitudes in at least 4 of the seven genes tested. These shifts involved delayed (*NpPRR5*, *NpLUX*) and advanced (*NpLHY* and *NpELF3*) maximum peaks of expression in seedlings from warmer environments (low altitude) (Fig. 6c-f). This environmental effect was not observed in *N. obliqua*, which showed synchronicity in maximum peaks of expression at both altitudes (Fig. 6g-j). Asynchronous seedlings of *N. pumilio* growing in warm environments showed reduced survival and accumulation of dry weight (Fig. 6k,m). This behavior was not evident in *N. obliqua* seedlings (Fig. 6l,n). These data are consistent with the idea that poor adjustment of the circadian clock is detrimental to the performance of the seedling in natural thermal environments.

**Fig. 6.**
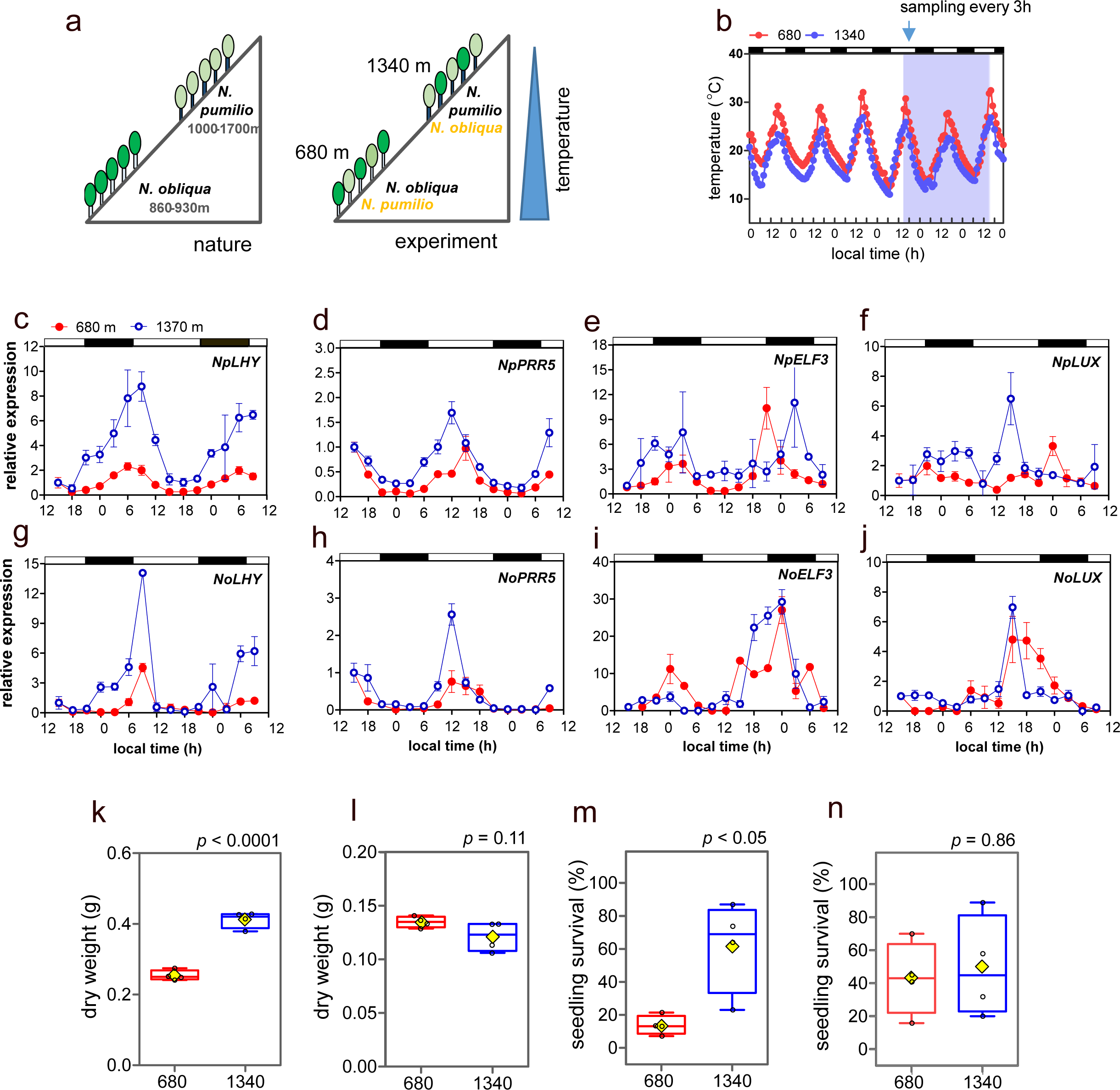
Expression of core oscillator genes and plant traits inside and outside the natural species’ thermal ranges. (a) Schematic representation of the natural distribution of the species in the forest (left) and in the experiment (right). (b) Air temperature at 680 and 1340 m a.s.l. Blue light area indicates the period of sampling. (c-f) Expression of *N. pumilio* clock genes *NpLHY*, *NpPRR5*, *NpELF3* and *NpLUX*. (g-j) Expression of *N. obliqua* clock genes *NoLHY*, *NoPRR5*, *NoELF3* and *NoLUX*. Data represent mean and SD of three technical replicates. Gene expression levels at time 0 was set to 1 and used as reference for all other time points. White and black rectangles represent day and night, respectively. Distribution of seedling dry weight and seedling survival of *N. pumilio* (k-m) and *N. obliqua* (l-n). Diamonds indicate the mean; box edges denote upper and lower quartiles; bars: whiskers show 10 and 90% percentiles; bold centre line indicates the median. Open circles indicate individual points that show the average of seedling mortality or dry weight in each plot (n =4 plots / species / altitude).

## Discussion

Until now, the effects of temperature on oscillator function to investigate thermal responses and susceptibility of plants have been relatively unexplored. Here, using two closely related tree species, we show that the upper thermal limits for accurate clock function are related to the thermal characteristics of their ecological niches, and to the ability of their clocks to match internal rhythms with cyclic changes in different natural thermic environments. The contribution of functional oscillators to an organism’s physiology and fitness has been extensively documented in different life kingdoms. A hallmark of such studies is that organisms with oscillators unable to match internal rhythms with external cyclic changes show reduced responses to the environment and lower fitness (7, 8, 27–29). Our findings, that seedlings of *N. pumilio* (high altitude species) had limited oscillator function in warmer (low altitude) zones of the forest, and reduced survival and growth is novel evidence that links disruption of oscillator function to poor tolerance of higher temperatures in the natural environment. By contrast, seedlings of *N. obliqua* (low altitude species) were able to maintain rhythms at higher temperatures than *N. pumilio*, and they showed similar survival and mortality in both environments of the temperature/altitude shift experiment, consistent with the fact that *N. obliqua* seedlings showed synchronised oscillators in both environments. Overall, our findings contribute to climate change biology, given that the ability of the oscillator to maintain rhythms at increasing temperatures is proposed to contribute to the capacity of plant species to respond to warming (30, 31).

The components of the core oscillator of the circadian clock are conserved from algae to angiosperms, both at the protein structural level as well as the gene expression level (time of day expression networks)(19, 32). This facilitates the identification of core oscillator orthologues among plant taxa and allows the extension of genetic studies related to circadian clock function to non-model plants. For instance, core clock genes of *N. pumilio* showed a high degree of conservation with their *Betula pendula*, *Populus trichocarpa* and *Arabidopsis thaliana* orthologues, and similar expression profiles (Figs. 2-5, Fig. S1-2, S4-9). Moreover, experiments under controlled conditions showed that time-of-day expression gene clusters were enriched in oscillating *Arabidopsis* orthologues at 20°C (Fig. 4) with an over-representation in biological processes previously reported to be under clock control^10^ (Tables S5-6). Our molecular approach also allowed us to determine that increasing temperatures affected not only the expression of core oscillator genes (Fig. 3, Fig. 5) but also of clock-regulated orthologues (Fig. 4). This suggests that temperature affects a wide range of physiological processes in seedlings growing under controlled conditions at least partially through an effect on oscillator function.

However, resolving the effect of temperature on oscillator function in complex natural environments is much more challenging compared to controlled environments, given the myriad factors that may be involved. Our multidisciplinary approach combined molecular, biochemical, bioinformatic and eco-physiological studies in controlled and natural environments, established a link between the sensitivity of the oscillator to increasing temperatures and the thermal characteristics of the species’ preferred thermal niche (Figs. 1-6). In addition, growth chamber experiments predicted biological processes that might be affected in seedlings growing in sub-optimal warm environments, since genes affected by heat included several clusters related to clock-controlled biological processes such as carbohydrate metabolism and growth(13, 33, 34). These processes relate to the phenotypes of seedlings that showed limited clock functioning in the warmer zones of the forest (Fig. 6k-n). A further point is that photocycles were apparently not able to override the effect of temperature on the oscillator of *Nothofagus* (Figs. 5-6). This finding is concordant with experiments in other plant species such as rice, where temperature played a more important role than photocycles in controlling the expression and phase of clock-related genes(36, 37). Similarly, temperature played a more important role than photoperiod in controlling gene expression in field-grown *Arabidopsis halleri* and *Brachypodium* plants (38, 39), and for setting the *Arabidopsis thaliana* oscillator under natural conditions (40). These results highlight the relevance of studying the effects of temperature on oscillator function as a focus area for assessing and comparing thermal responses of plants.

Nevertheless, at the plant species level, it is important that studies exploring responses to global warming consider other factors in addition to the role of the circadian clock. For instance, in *Nothofagus*, seed dormancy responses to temperature determined temporal patterns of germination across altitude (41). Additionally, *N. pumilio* seedlings emerging in the local soil environment showed reduced leaf area in warmer zones of the forests, even though differences in mortality between altitudes were not evident under those experimental conditions (42). Probably, other factors such as biotic interactions (43–45) and soil properties (46) influence seedling recruitment in the wild, and this may or may not be independent of clock function (47–49). Furthermore, this study focusses on the effect of temperature on just one aspect of forest dynamics – namely, seedling growth. Other aspects of forest dynamics have recently been assessed for their thermal susceptibility, such as spring phenology (3, 4, 50) and leaf senescence (51) of adult trees. Interestingly, both of these traits proved to be controlled, at least in part, by the oscillator (52, 53). Further studies are required on other traits at different stages of tree growth under natural conditions to investigate the role of the circadian clock and the potential effect of temperature on its performance. Despite these caveats, our work represents the first empirical study in testing the effect of temperature on circadian clocks as a proxy for analysing thermal vulnerability in plants. Our results also highlight the potential role of a resonating oscillator in ecological adaptation as well as in forest regeneration in a warming environment. Indeed, the effect of increasing temperatures on oscillator function may be one factor which constrains the regeneration of the dominant species *N. pumilio*, potentially jeopardising the integrity of the ecosystem of the Andean-Patagonian forests.

Finally, it is important to note that artificial selection on circadian clock genes affecting photoperiod sensing throughout the history of agriculture has allowed the geographic expansion of several cultivated plant species to extreme latitudinal ranges (54–57). In this context, the investigation of genes responsible for thermal stability of the circadian clock may contribute to the selection of plant genotypes/populations with increased resistance to warming. These would be of great value for forest conservation and crop production in the face of the threat imposed by global climate change.

## Materials and Methods

### Site and species’ descriptions

Site and species characteristics were described by Arana et al. 2016 (58), and in *SI Materials and Methods*. Experiments were carried out in an old-growth mixed temperate forest in Lanin National Park, Neuquén province, Argentina (latitude: 40°08’20’’ S, longitude: 71°28’41’’ W, mean annual precipitation: 2100 mm.y^-1^). We performed all sampling and experiments in two enclosed permanent plots at 680 and 1340 meters above sea level (m.a.s.l), that is, at the natural distribution area of *Nothofagus obliqua* and *N. pumilio* respectively. We chose homogeneous forest habitats for field experiments, representative of old-growth stands and characterized by a canopy and understory cover that yields Red : Far Red ratios ranging from 0.5 to 0.7 at the soil surface. These sites showed clear differences in air temperatures, with the lower site exhibiting a warmer profile than the higher sites, and similar drought index values along the seasons (Fig. 1). Climatic data recording and processing is described in *SI Materials and methods*.

### Identification and structural characterization of circadian clock genes in *N. pumilio*

RNA-seq reads of *N. pumilio* (59) were merged and aligned to the draft genome of the species with STAR (60) to create a reference-based transcriptome using Scallop (61). Peptides were predicted with Transdecoder v5.5.0 (62). The predicted *N. pumilio* proteome was aligned using blastp (63) v2.11.0 against the reference annotated proteomes of *Arabidopsis thaliana* (64), *Populus trichocarpa* v3.0 (65) and *Betula pendula* (66) (parameters: -max_target_seqs 1 - outfmt 6 -evalue 1e-5). For each proteome, protein domains were searched with hmmer (67) v3.2.1 using the Pfam (68) v34.0 database (Table S1).

Circadian clock genes were classified in two groups: (1) core oscillator genes (Fig. 2) and (2) genes with other functions in addition to circadian regulation (Fig. S1) according to previous work (69, 70). Genes of interest were individually searched in the reference proteomes and for those where Pfam domains were not found, disordered residues along the protein sequences were predicted using flDPnn (71) vDecember 2021. Protein domain and disordered regions were plotted and exported using DoMosaics (72), and edited in Inkscape v1.1.2 (Table S10, Figures 2, S1 and S2). Details of phylogenetic analysis are provided in *SI Extended Methods*.

### Seed sources

Seed sources are described in *SI Extended Methods*.

### Experimental conditions for determining the effect of temperature on clock functioning in controlled environments

Experiments were performed in growth chambers (Percival Scientific, LT-36VL, Iowa, USA). To study the effect of warm temperatures (34°C) in the expression of core oscillator genes of *N. pumilio* under circadian conditions, 1 year-old plants were exposed to 12 h light, irradiance: 200 μmol m^−2^ s^−1^ / 12 h dark (LD12/12) at 20 °C followed by continuous light, irradiance: 100 μmol m^−2^ s^−1^ (LL), at either high (34°C) or control (20°C) temperature treatments. The onset of the free-running conditions is known in circadian literature as ZT0 (Zeitgeber). Samples were collected every three hours starting at ZT24 (Fig. 3A). Each sample consisted of a pool of one whole leaf from 10 different seedlings. Samples were immediately frozen in liquid nitrogen and stored at -80°C until the RNA extraction.

For experiments in light/dark photocycles, *N. pumilio* and *N. obliqua* plants were exposed to LD12/12 at 20 °C for 7 d and then to LD12/12 at either 20, 31 or 34 °C. Samples were collected every three hours starting 24 hours after the release into the different temperatures, pooling leaves from 10 different seedlings; (Fig. 4A). Strategy for setting the temperatures for inter-specific comparisons is described in *SI Extended Methods*.

### Primer design

Primer sequences (Table S11) were designed using Primer3 (www.primer3.org). Primer design is described in detail in *SI Extended Methods*.

### Quantification of gene expression

RNA extraction, cDNA synthesis and quantitative PCR were performed as described by Estravis-Barcala et al. 2021 (59), Gene expression graphs were obtained using GraphPad Prism v5 (https://www.graphpad.com/features).

### Effect of temperature on global patterns of gene expression under constant conditions

To investigate changes in global patterns of gene expression between different moments of the day and temperatures in *N. pumilio*, we analysed the RNA-seq datasets described in Estravis-Barcala et. al 2021 (59). This data consisted of four different conditions: subjective dawn at 20°C, subjective dusk at 20°C, subjective dawn at 34°C, subjective dusk at 34°C. We obtained the gene read counts for each biological replicate after mapping their corresponding RNA-seq to the *N. pumilio* draft genome using Star (60) (parameter –alignIntronMax 25000). We contrasted the global patterns of gene expression between subjective dusk and dawn at both temperatures with DESeq2 (73). For enrichment analysis, phase assignation and evaluation of orthologues of direct clock targets we used the databases described in Nagel et al. 2015 (74), Gendron et al. 2016(26) and Harmer et al. 2000(13), the last one available in DIURNAL (75) (LL12_LDHH acronym) http://diurnal.mocklerlab.org. <u>Data analysis are</u> described in *SI Materials and Methods*.

### Field experiments

Seedlings from both species were germinated as indicated in *SI Materials and Methods*, and grown separated by at least 5 cm in 2,200 cm^3^ with soil from the IFAB campus. Seedlings were grown in the greenhouse for 3 months with daily water sprinkler irrigation before taking them to the field in late spring (mid-December 2012). Seedlings were transplanted in the forest in the original soil of the pots, to avoid any bias due to soil differences between altitudes. Each planted pot constituted a plot of the experiment, and seedlings were transplanted in the field in four plots per species per altitude. One week after transplantation, we scored the initial number of seedlings for the experiment, totaling 320 seedlings for *N. obliqua* and 362 seedlings for *N. pumilio*. These seedlings were sampled after one month of acclimation in the forest, in summer (mid-January 2013). Samples were collected in synchronic way at both altitudes, every three hours. Each sample consisted in a pool of five leaves, coming from different seedlings (collecting one leave/seedling). Seedlings were chosen at random in the different plots, referenced using plastic sticks, and harvested just once. Leaves were immediately frozen in liquid nitrogen, taken to the lab and kept at -80°C until RNA extraction. Estimation of plant survival, dry weight and statistical analysis are detailed in the section *SI materials and methods*.

## Acknowledgements

This work was supported by the Agencia Nacional de Promoción de la Investigación, el Desarrollo Tecnológico y la Innovación, Argentina (PICT 2011/2250, 2017/2656, 2020/02146), CONICET, Argentina (PIP 2020-11220200102254CO) and INTA, Argentina (2019-PD-E6-I116 and 2023-PD-L01-I085), and Instituto Milenio iBio ICN17_022– Iniciativa Científica ANID, Chile and ANID – Millennium Science Initiative Program – ICN2021_044, Chile. Maximiliano Estravis-Barcala was supported by CONICET doctoral and postdoctoral fellowships. We thank Dr Peter Chandler (CSIRO, Australia) for helpful comments on the manuscript and English style. We are eternally thankful to Dr. Todd Mockler (Donald Danforth Plant Science Center) for the creation of the diurnal webpage and to Dr. Todd Michael (The Salk Institute for Biological Studies) for the provision of the LL12_LDHH and LD12HH_ST diurnal databases. We thank Dr Alejandro Aparicio, Carolina Soliani, Paula Marchelli, Jorge Bossi, Alejandro González for their help in the field sampling for gene expression analysis, and Nahuel Huapi and Lanin National Parks for permissions on seed collection and field experiments. We are grateful to Marcelo González Peñalba, Liliana Lozano, Martín Lara and Carlos Clericci (Lanin National Park) for their support and advice during experiments in the altitudinal gradient. We thank Dr Alvaro Gutiérrez (University of Chile) and Fernando Umaña (INTA EEA Bariloche) for the provision of the shape files of the distribution of *N. obliqua* and *N. pumilio* along the eastern and western Andes, Diego Schell (CONAE) for the confection of the species distribution maps and Dr Verónica Pancotto (CADIC, Tierra del Fuego, Argentina) for the *N. pumilio* forests at high latitudes of picture in Fig. 1.

