## Extended Methods and Supplementary Figures for "Effect of temperature on circadian clock functioning of trees in the context of global warming"

#### Site descriptions

Experiments were carried out in an old-growth mixed temperate forest in Lanin National Park, Neuquén province, Argentina (latitude: 40° 08’ 20’’ S, longitude: 71° 28’ 41’’ W). In this forest, *Nothofagus obliqua* is the dominant tree species in the lower elevations ca. 650 meters above sea level (m.a.s.l.) up to ca. 930 m.a.s.l. In contrast, the highest elevations of the forest are dominated by *N. pumilio*, from ca. 1000 m.a.s.l. to ca. 1700 m.a.s.l. The site is located in the humid region of the eastern Andes with a mean annual precipitation of 2100 millimetres per year (mm.y$^{-1}$). We performed all sampling and experiments in two enclosed permanent plots at 680 and 1340 m.a.s.l, that is, at the natural distribution area of *N. obliqua* and *N. pumilio* respectively. We chose homogeneous forest habitats for field experiments, representative of old-growth stands and characterized by a canopy and understory cover that yields Red : Far Red ratios ranging from 0.5 to 0.7 at the soil surface. These sites showed clear differences in air temperatures, with the lower site exhibiting a warmer profile than the higher sites, and similar drought index values along the seasons (Fig. 1). In both sites, understorey vegetation is dominated by *Chusquea culeou* Devaux.

#### Species’ descriptions

*Nothofagus* is the only genus in the monotypic family Nothofagaceae, with 35 extant species in the southern hemisphere. In modern times, *Nothofagus* has a disjoint distribution in Oceania and southern South America. A large fossil record, mostly leaves and pollen, can be found throughout the ancient Gondwanic province of Weddell, which has made *Nothofagus* a key genus for understanding the origin, migration, diversification, and change of patterns in the southern biota[1,2]. Particularly, *Nothofagus pumilio* is one of the most widely distributed species of the Patagonian forests, and it inhabits an iconic latitudinal gradient, from the northern Patagonian Andes and central Chilean region (35˚S) to the high latitudes at Tierra del Fuego (55˚S). This southern region of the Andes hosts rainforest and sub-Antarctic temperate forests that embrace an extraordinary ecological diversity across different environments[3,4] that will be affected by increasing temperatures according to predictions of global climate change[5]. *N. pumilio* shows an unusual dependence of its altitudinal distribution with latitude not found in other native species of the region. It ranges in elevation from 0 to 2000 m.a.s.l., but north of latitude 41˚S it grows only in the sub-Alpine colder zone where it commonly forms the treeline. On the other hand, in the southern part of its range, in colder environments of high latitudes, it occurs both at high (treeline) and low (sea level) elevations[6]. This suggests that the species lacks adaptation to warmer environments. In contrast, *N. obliqua* has a more restricted latitudinal distribution, from latitude 34 to 41˚S in Chile, and from latitude 36 to 40˚S in Argentina, mostly in lake basins of Lanin National Park[6,7]. *N. obliqua* coexists with *N. pumilio* in northern Patagonia, but it occurs in lower altitudes than *N. pumilio*. The region of this study, the Lacar lake basin of Lanin National Park around latitude 40°S and longitude 72˚S, features areas of *N. pumilio* and *N. obliqua* forests. Whereas the latter is ubiquitous at lower altitudes, between 600 and 800 m.a.s.l, *N. pumilio* thrives from 1000 m.a.s.l. up to the treeline, which at this latitude sits at approximately 1700 m.a.s.l.[6,8] (Fig. 1).

**Climatic data**

Air temperature and moisture were monitored using Onset HOBO® data loggers (http://www.onsetcomp.com/). Climatic data were processed using R[9]. Red : Far Red ratios at the soil surface of the experiments were measured near midday (12:00–15:00 h) at all sampling dates using a 660/730 nm quantum sensor (Skye Instruments Ltd, Powys, UK).

Drought index was calculated according to Arana et al 2016[10], using the following equation:

$$((3 * Tmax) + Tmin) / (1 + HRm)$$

where Tmax is the daily maximum temperature, Tmin is the daily minimum temperature and HRm is the daily average air humidity.

**Seed sources**

All the experiments were performed using seedlings produced from seeds collected from natural *N. pumilio* and *N. obliqua* populations. For field experiments involving gene expression, quantification of seedling mortality, and dry weight, *N. pumilio* plants were produced using seeds collected during 2012 from Challhuaco (latitude: 41º15'29" S, longitude: 71º17'07" W, mean annual precipitation: 1000 mm.$y^{-1}$, average altitude: 1175 m.a.s.l), Chapelco (latitude: 40º11'59" S, longitude: 71º20'08" W, mean annual precipitation: 1500 mm.$y^{-1}$, average altitude: 1204 m.a.s.l.) and Cerro Colorado (latitude: 40º08'59" S, longitude: 71º23'15" W, mean annual precipitation: 1130 mm.$y^{-1}$, average altitude: 1400 m.a.s.l.). *N. obliqua* plants were produced using seeds collected during 2012 from Quila-Quina (latitude: 40º09'07" S, longitude: 71º26'37" W, mean annual precipitation: 1800 mm. $y^{-1}$, average altitude: 670 m.a.s.l.) and Yuco (latitude: 40º09'07" S, longitude: 71º30'39" W, mean annual precipitation: 2100 mm. $y^{-1}$, average altitude: 680 m.a.s.l.). Chamber experiments were performed using *N. pumilio* plants produced from seeds collected from Challhuaco and *N. obliqua* plants produced using seeds collected in Quila-Quina during 2012 and 2015. These constitute representative forests of the two species, that produce relatively good amounts of seeds along the different years. This property is relevant to ensure the availability of enough amounts of seeds to explore thermal ranges for proper oscillator function at different conditions along the years of experimentation (i.e. under free running and photocycles), given the seed-masting habit of the genus[11].

In all the cases we harvested seeds from at least 25 individual trees located at a minimum distance of 30 m in order to preclude family relationships. Equal amount of seeds from each mother plant were pooled for the experiments. For field experiments that used different seed origins, equal amount of seeds from each population and mother plant were pooled to ensure equal representation of origins and mother plants in the experiments.

**Phylogenetic analysis**

To further support the orthology of the newly described *N. pumilio* clock genes, phylogenetic analyses were performed for those genes which were measured by qPCR. Peptidic sequences of clock-related members of the PRR and MYB gene families from *N. pumilio*, *A. thaliana*, *B. pendula* and *P. trichocarpa* were aligned using MUSCLE version 5.1[12]. After alignment, the best molecular evolution model for each matrix was searched by ProtTest version 3.4.2 with default parameters[13]. Maximum likelihood phylogenetic reconstructions were performed by RAxML version 8.2.12 with 10,000 bootstrap replicates[14]. *A. thaliana* orthologs of closely related genes (AtPRR9 for the PRR family and AtRVE7 for the MYB family) were used as outgroups. Phylogenetic trees were plotted, edited, and exported using EMBL's interactive Tree of Life (iTOL) tool[15]. In the case of ELF3 and GI, no reconstruction was possible since they are not members of multigenic families, and they don't have related proteins by sequence.

**Primer design**

Primers were designed in regions encompassing *N. pumilio* exons with homology to *A. thaliana*. Each primer pair was verified *in silico*, confirming that it had a unique amplification site in the draft genome (Table S11). Alignment against the draft genome was visualised with the Integrative Genome Viewer IGV (www.igv.org; Fig. S3). Choice criteria for DER2.2 as housekeeping gene is detailed in the *SI Materials and Methods*. All primer pairs designed following this protocol successfully amplified single PCR products in both *N. pumilio* and *N. obliqua*.

### Choice of DER2.2 as housekeeping gene

The reference gene, DER2.2 (*Arabidopsis thaliana* orthologue ID: At4g04860) was chosen since its expression didn't change in response to neither temperature nor time of the day in RNA-seq experiments of *N. pumilio*[16] (DESeq2 *p*-value > 0.95 for all pairwise comparisons between time of day and temperature treatments performed in this study; see differential expression protocol in Materials and Methods). Additionally, in *A. thaliana*, DER2.2 orthologue was classified as (1) not-cycling in DIURNAL[17] database (available in http://diurnal.mocklerlab.org) in conditions similar to those of our experiment: entrainment in light/dark 12h/12h cycles, followed by continuous light (LL12_LDHH in DIURNAL acronym, complete experiment described in Harmer et al. 2000[18]) and (2) not affected by heat, as its expression kept constant under a 3-hour-long 38°C treatment[19]. Both in *N. pumilio* and *A. thaliana* DER2.2 showed a mid-to-high expression (DESeq2 base mean in *N. pumilio* experiments > 20 for all pairwise comparisons; DIURNAL expression data in *A. thaliana* varied between the median and the third quartile of expression for all loci).

### Seedling production and nursing

We produced plants in the greenhouse, using the following protocol. After 60 days of stratification at 4°C in 9 cm-diameter Petri dishes with cotton moistened with 5ml of a water solution with 1% of the fungicide Vitavax-Flo (Lujan Agrícola, Argentina), the seeds were sowed in trays until germination and cotyledon emergence occurred. At this moment, seedlings were transplanted to individual, 265 $cm^3$ pots arranged in 28-pots plastic trays inside of a temperature- and irrigation-controlled greenhouse at IFAB, San Carlos de Bariloche, Argentina. These pots contained an inert mixture of equal parts of volcanic sand and peat. During one season, seedlings were subjected to a greenhouse growing protocol developed for *Nothofagus* species[20]. Briefly, irrigation was conducted to saturation (water dripping from the bottom of the trays). Macronutrients in the form of NPK (Nitrogen-Phosphorus-Potassium) were provided through sprinklers on top of the plant trays. On the other hand, micronutrients (Mn, Zn, Fe, Cu, Cl, B, Mo) were supplied via foliar application during the maximum growth stage. The growing cycle consisted of three stages: establishment (root development; NPK formulation: 10-45-16), maximum growth (leaf and stem development; NPK formulations: 18-7-17 and 14-0-14 formulation, with a weekly alternation between formulations), and hardening (bud and secondary stem development; NPK formulation: 4-27-38). The total length of the growing cycle is 7 months (beginning of September to end of March in the southern hemisphere). Together with the stratification and germination phases, this protocol produces 60 cm tall plants in 9 months. Once the hardening stage was over and the plants dropped their leaves, they were kept in the greenhouse to be used in growth chamber experiments from the next summer.

**Effect of temperature on global patterns of gene expression under constant conditions**

After read counting and differential expression analyses, we sought to assess whether the differentially expressed transcripts between the subjective dawn and dusk were enriched in cyclic *Arabidopsis thaliana* orthologues. For this, we first selected those significant expressed transcripts ($p$-value < 0.05). We then obtained their corresponding *A. thaliana* orthologues (see "Identification and structural characterization of circadian clock genes in *N. pumilio*" in Materials and Methods) and removed common IDs. We retrieved the phases (hour of maximum expression) and correlation to cycling models[18] from the aforementioned DIURNAL database LL12_LDHH. This database is the one that better fits with our experimental conditions: entrainment with photocycles of 12 hours light / 12 hours darkness and sampling in continuous light at 22˚C. See Harmer et al. 2000[18] for detailed experimental conditions. We contrasted the distribution of correlations of our datasets against the distribution of the whole *A. thaliana* IDs in LL12_LDHH. We considered that a specific dataset was enriched in genes with cyclic patterns of expression when the observed correlation distribution of the oscillating genes of the experiment (MBPMA cut-off of 0.8 as described in Nagel et al. 2015[21]) was significantly greater than the expected correlation distribution of the DIURNAL database LL12_LDHH (Kolmogorov-Smirnov $p$-value < 0.05). As proof of concept, we applied this methodology in the set of direct CCA1 targets genes described in Nagel et al. 2015[21] (Fig. S12). Once we assessed the enrichment of circadian genes in a particular dataset, the question that raised was whether such genes had enrichment in specific phases. For this purpose, we compared the observed genes in each phase against a uniform distribution calculating the log2 Fold Change in each phase (Fig. 4e-f). Gene Ontology (GO) term enrichment analyses were performed using the GO platform[22] available in https://geneontology.org/. We analysed the *A. thaliana* orthologues of cycling genes differentially expressed between subjective dawn and dusk at 20˚C, 34˚C and those exclusively regulated at 20˚C, 34˚C, or commonly regulated at both temperatures.

To study the effect of disruption of clock functioning by warm temperatures exclusively on orthologues of direct clock targets, we mapped the genes differentially regulated at 20˚C against *Arabidopsis* orthologues of direct TOC1 targets described in Gendron et al. 2012[23] and analyzed their expression at 34˚C. We chose this dataset of genes because (1) it contains targets identified by direct interaction between TOC1 and DNA regions through Chip-seq, and (2) considering the works available in the literature that investigate direct clock targets in constant light conditions (LL) through Chip-seq[21,23–29], the data set published by Gendron et al. 2012[23] constitutes the largest gene list of direct clock targets. We found a total of 1733 orthologues of TOC1 direct clock targets in the transcriptome of *N. pumilio*, where 1051 of those genes constituted unique IDs. These 1051 genes represent 60% of the genes of the original published list. Using all the identified orthologues of direct TOC1 targets (n=1733), we then searched for those genes that were differentially regulated between moments of the day at 20˚C, to evaluate the effect of heat (34˚C) in their regulation. For this purpose, we searched for those genes that (1) were differentially expressed between times of the day at 20˚C (2) showed consistence between the two biological replicates; this means that the two biological replicates at dawn showed a higher expression than the two biological replicates at dusk, or vice versa. The latter is an important point given that DESeq2 reports the differential expression results considering only the average of the different replicates. We found that 15.35% (n=265) of the orthologues met this criterion, and this in concordance with the fact that only a group of genes (30%) of the original *Arabidopsis* list showed a cyclic behaviour (correlation > 0.8) in the Diurnal database LD12_LLHH. We applied hierarchical clustering with pheatmap (https://datasciencetut.com/the-pheatmap-function-in-r/) to visualise the regulation of this group of *N. pumilio* genes, considering all the biological replicates of the treatment at 20 and 34˚C.

**Experimental conditions for determining the effect of temperature on clock functioning in controlled environments**

We first investigated the effect of warm temperatures (34°C) in the expression of core oscillator genes under circadian conditions, also known as free-running conditions. For this purpose, we incubated 1 year-old plants for 10 days in growth chambers (Percival Scientific, LT-36VL, Iowa, USA) at 20°C with 12 hours light (200 μmol $m^{-2}$ $s^{-1}$) and 12 hours darkness at 20°C. The exposure of plants to cyclic conditions, in this case photocycles, allows the oscillator to be fully entrained by environmental conditions before the analysis of its performance at control or high temperatures. Then, the plants were subjected to continuous light (100 μmol

$m^{-2}$ $s^{-1}$), so called circadian conditions, and to either a high (34°C) or control (20°C) temperature treatment. In this second step, irradiance is reduced to 50% to keep the daily amount of photons received by the plants constant during all the experiment. Additionally, 100 µmol $m^{-2}$ $s^{-1}$ represents moderate photosynthetic photon flux densities for the species[4] and therefore reduces the possibility of high light irradiance-mediated stress during prolonged periods of exposure to light in the free-running condition stage. The onset of the free-running conditions is known in circadian literature as ZT0 (Zeitgeber). Samples were collected every three hours starting at ZT24 (Fig. 3A). This timing of sampling allows the stabilization of the oscillator during the first 24 hours after the release into continuous light. Each sample consisted of a pool of one whole leaf from 10 different seedlings. Samples were immediately frozen in liquid nitrogen and stored at -80°C until the RNA extraction.

For experiments in light/dark photocycles (also called diurnal conditions), the procedure was the same, except that the light/dark cycles (200 µmol $m^{-2}$ $s^{-1}$) were continued after the entrainment phase, when setting the temperature treatment. In order to compare the effect of the temperature in the performance of the oscillator of *N. pumilio* and *N. obliqua*, we analysed the expression of core genes at 31°C and 34°C to study potential inter-specific differences in the effect of temperature on circadian clock functioning. For *N. pumilio*, a temperature treatment of 28°C had been first performed, where we found a gene expression pattern similar to the control temperature at 20°C (Fig. S11, Fig. 5f, Fig. S7d). Samples were collected in the same way as for circadian experiments (every three hours starting 24 hours after the release into the different temperatures, and pooling leaves from 10 different seedlings; Fig. 4A). Each experiment (both circadian and diurnal, at each specific temperature) was performed twice in the same growth chambers, using different seedlings.

**Estimation of plant survival and dry weight**

Plant survival and dry weight were scored at the end of the growth season in intact seedlings. The percent of survival was estimated per plot (each plot = one replicate, yielding n=4 for each species and altitude) according to the following equation:

% survival = (final number of seedlings / initial number of seedlings) * 100

Survival experiments involved a total of 160 seedlings for *N. obliqua* and 202 seedlings for *N. pumilio*.

For dry weight, surviving seedlings were taken to the laboratory, and dried in a stove until constant weight. Dry weight data was calculated as the average of the dry weight of the seedlings per plot (each plot = one replicate, yielding n=4 for each species and altitude), totaling the analysis of 73 seedlings for *N. obliqua* and 69 seedlings for *N. pumilio.*

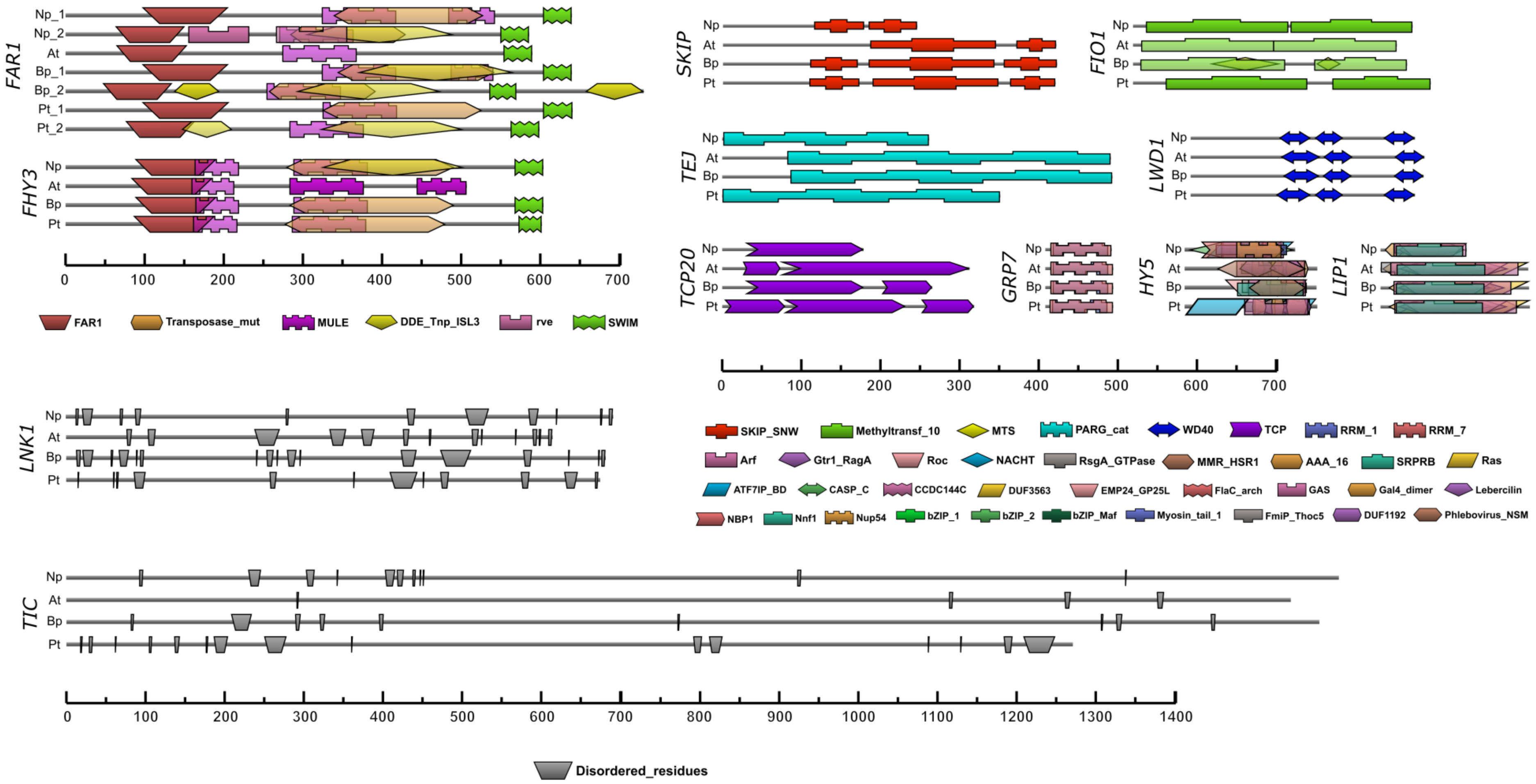


Fig. S1 | **In silico characterization of circadian clock genes with other functions in addition to circadian regulation**. Pfam protein domains were predicted using HMMER. Positions and p values of domains are listed in Table S10. In proteins lacking predicted Pfam domains (TIC and LNK1), disordered regions were predicted using flDPnn. The scale bar represents the amino acid number. Np: *Nothofagus pumilio*. At: *Arabidopsis thaliana.* Bp: *Betula pendula*. Pt: *Populus trichocarpa*.

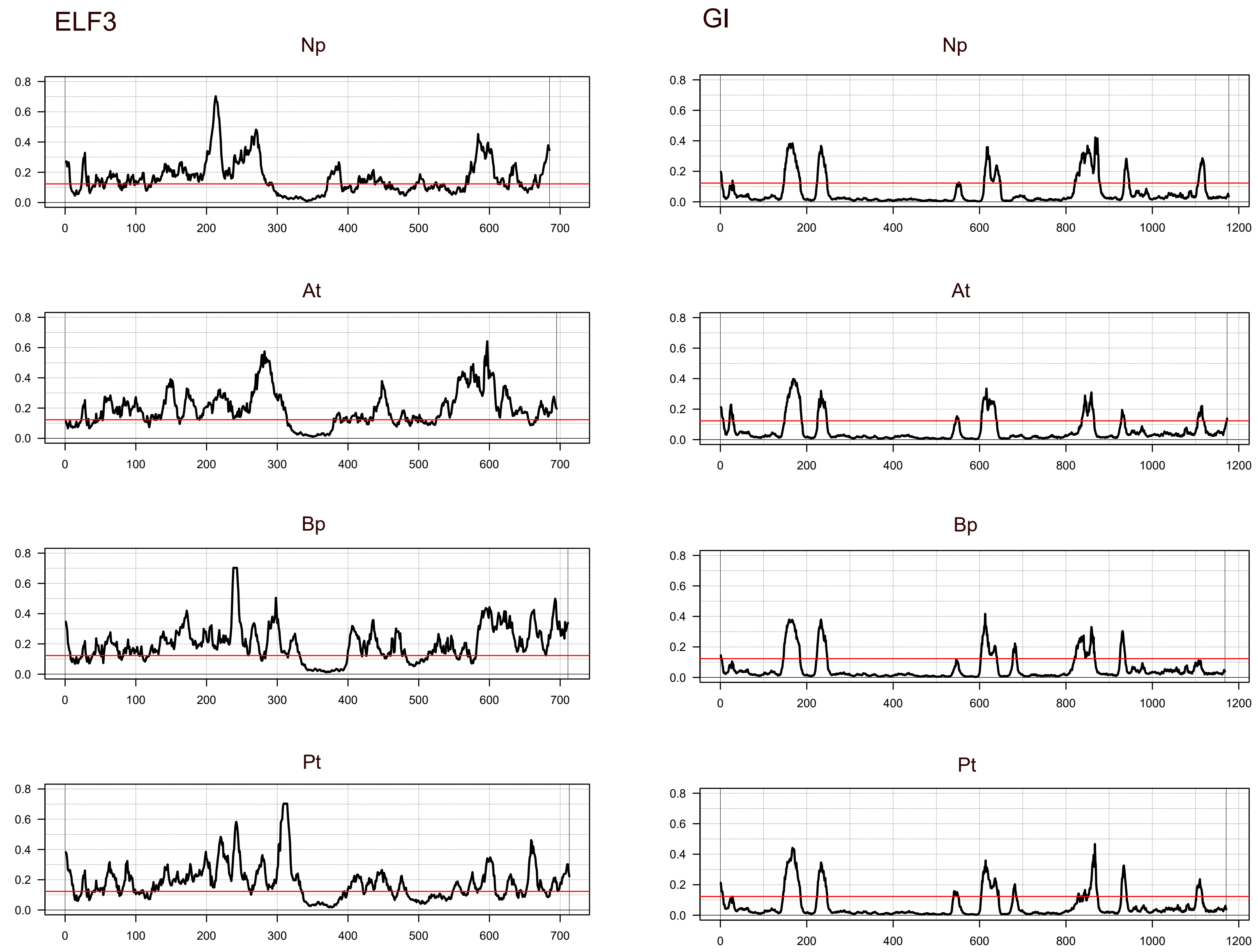


Fig. S2 | **Disorder score along amino-acid sequences of EARLY FLOWERING 3 (ELF3) and GIGANTEA (GI) proteins**. Disorder score per residue was calculated using the flDPnn algorithm. The red line indicates the disorder threshold (0.123) used to infer disorder regions according to Parra et al. 2023 (doi: 10.1002/yea.3853). Regions over the threshold are referred to as intrinsically disordered regions (IDRs). Np: *Nothofagus pumilio*. At: *Arabidopsis thaliana*. Bp: *Betula pendula*. Pt: *Populus trichocarpa*. Red line indicates cut-off to determine disordered regions.

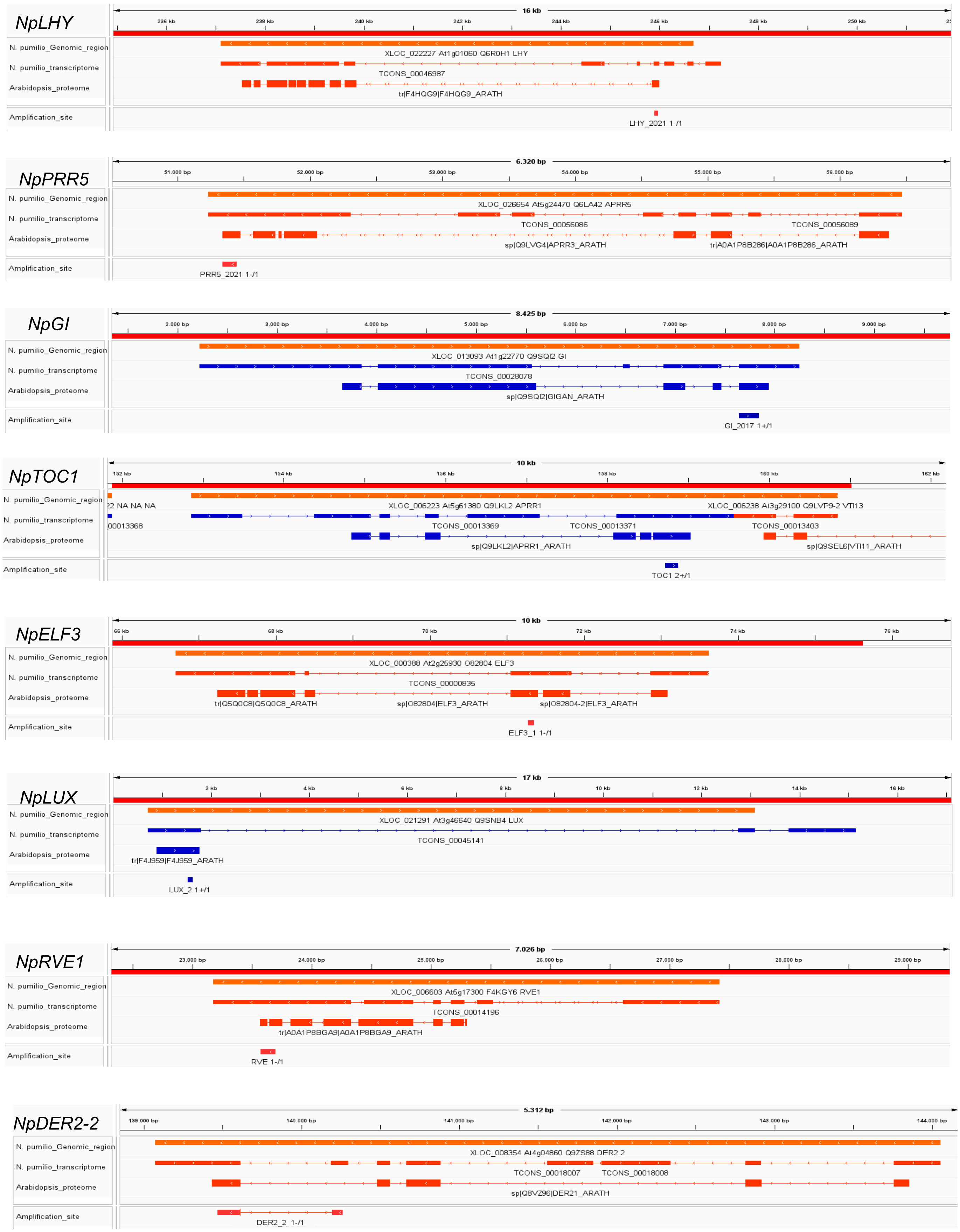


Fig. S3 | **IGV visualization of amplification sites of genes chosen for this study**. Primers sequences were mapped to the alignment of the *N. pumilio* transcriptome and the *Arabidopsis thaliana* proteome to the *N. pumilio* draft genome. Positions in the genomic scaffold are indicated with numbers in the upper red line. Gene locus is indicated in orange. Blue and red colours indicate sense and anti-sense alignment, respectively. Amplification sites correspond to the *N. pumilio* transcriptomic region flanked by the primers. Annotations are described in Table S1. Briefly, XLOC_number denotes the locus in the *N. pumilio* draft genome, which is followed by the corresponding *A. thaliana* genomic ID, Uniprot code, and abbreviated gene name. TCONS_number corresponds to the annotation of the *N. pumilio* transcript. *A. thaliana* annotations correspond to Uniprot codes.

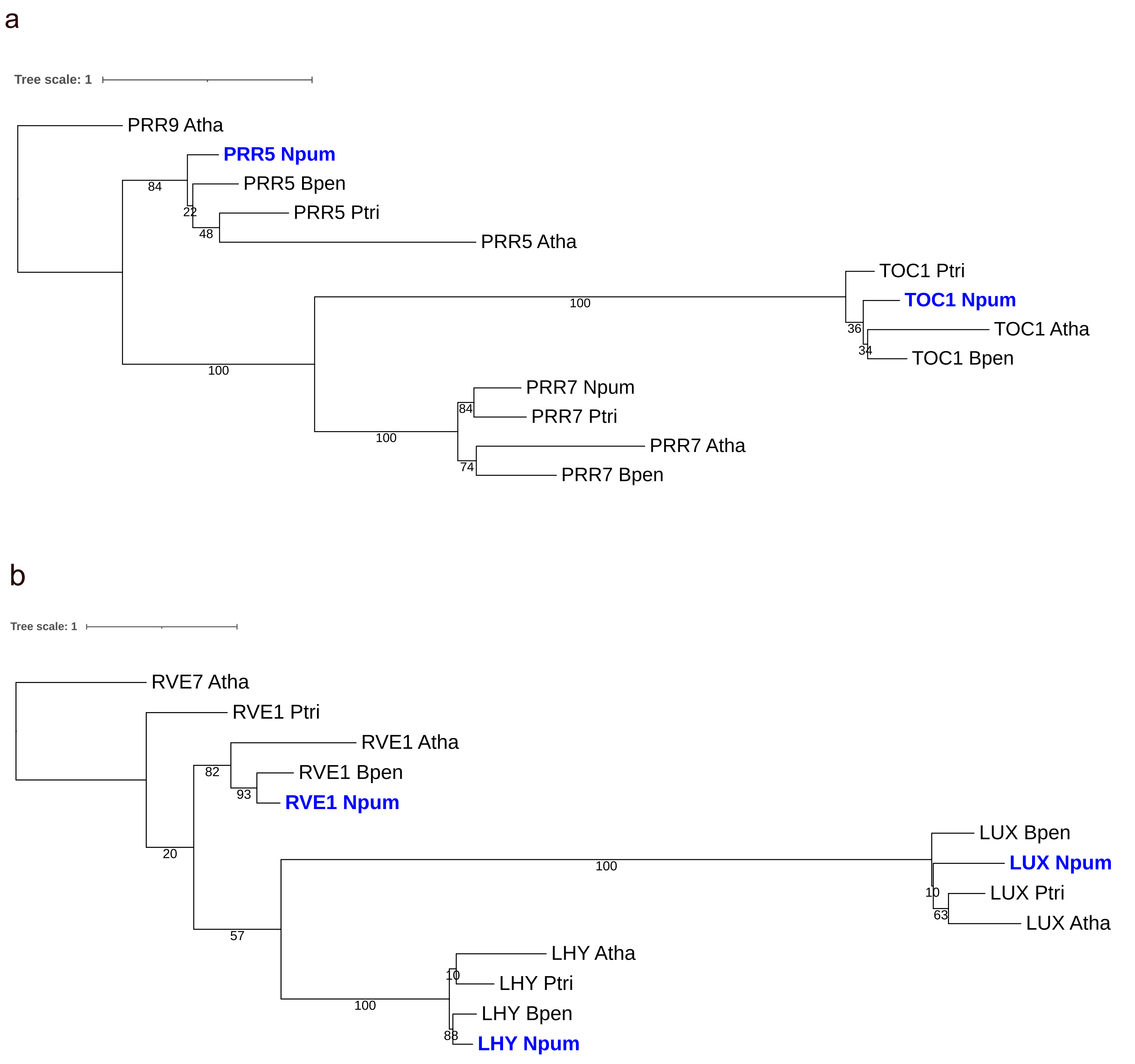


Fig. S4 | **Maximum Likelihood phylogenetic trees for genes measured by qPCR**. (a) PRR family. (B) MYB family. Bootstrap support is shown in each bipartition. Branch length corresponds to the mean number of substitutions per site. Bold blue tips indicate genes measured by qPCR in this study. Npum: *Nothofagus pumilio*. Atha: *Arabidopsis thaliana*. Bpen: *Betula pendula*. Ptri: *Populus trichocarpa*.

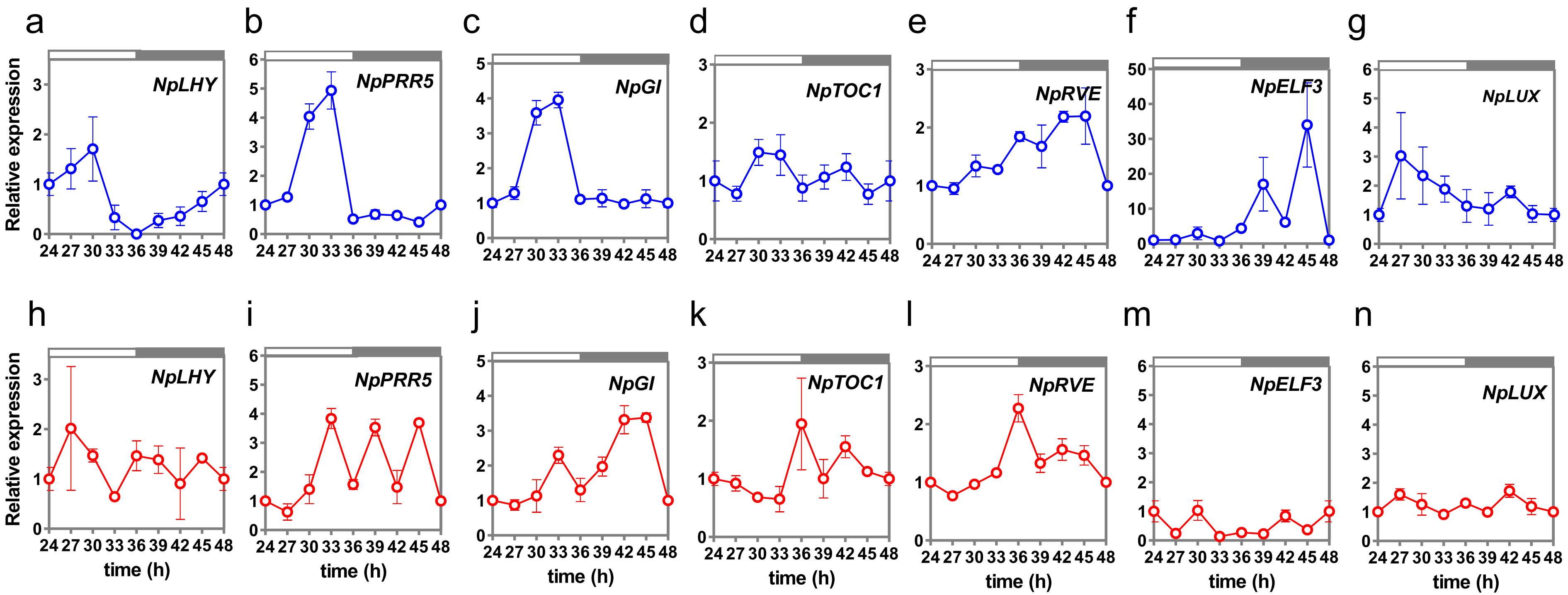


Fig. S5 | **Influence of temperature on the expression of clock genes under circadian conditions.** Biological replicate of experiment shown in Fig. 3. Seedlings were entrained with 12h light / 12h darkness photocycles at 20°C and then exposed to continuous light (free-running conditions) at 20°C or 34°C. Samples were taken every 3h during the second day after release into continuous light. (b-h) Expression of *NpLHY, NpPRR5*, *NpGI*, *NpTOC1*, *NpELF3*, *NpLUX* and *NpRVE1* at 20°C (blue symbols) or 34°C (red symbols). Data represent mean and SD of three (20°C) or two (34°C) technical replicates. Gene expression levels at ZT0 were set to 1 and used as reference for all other time points.

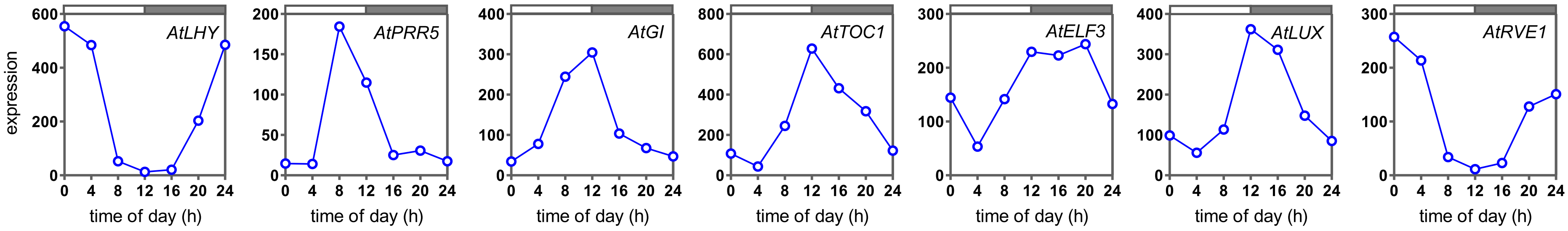


Fig. S6 | **Transcript levels of *Arabidopsis thaliana* clock genes *AtLHY*, *AtPRR5, AtGI*, *AtTOC1*, *AtELF3*, *AtLUX* and *AtRVE1* under circadian conditions**. *A. thaliana* seedlings were entrained with photocycles (12h light / 12h darkness) at 22°C, and then exposed to continuous light. Sampling was performed every 4h under continuous light. Data correspond to microarray experiments taken from DIURNAL (http://diurnal.mocklerlab.org/, LL12_LDHH database). The complete experiment is described in Harmer et al. 2000 (doi: 10.1126/science.290.5499.2110).

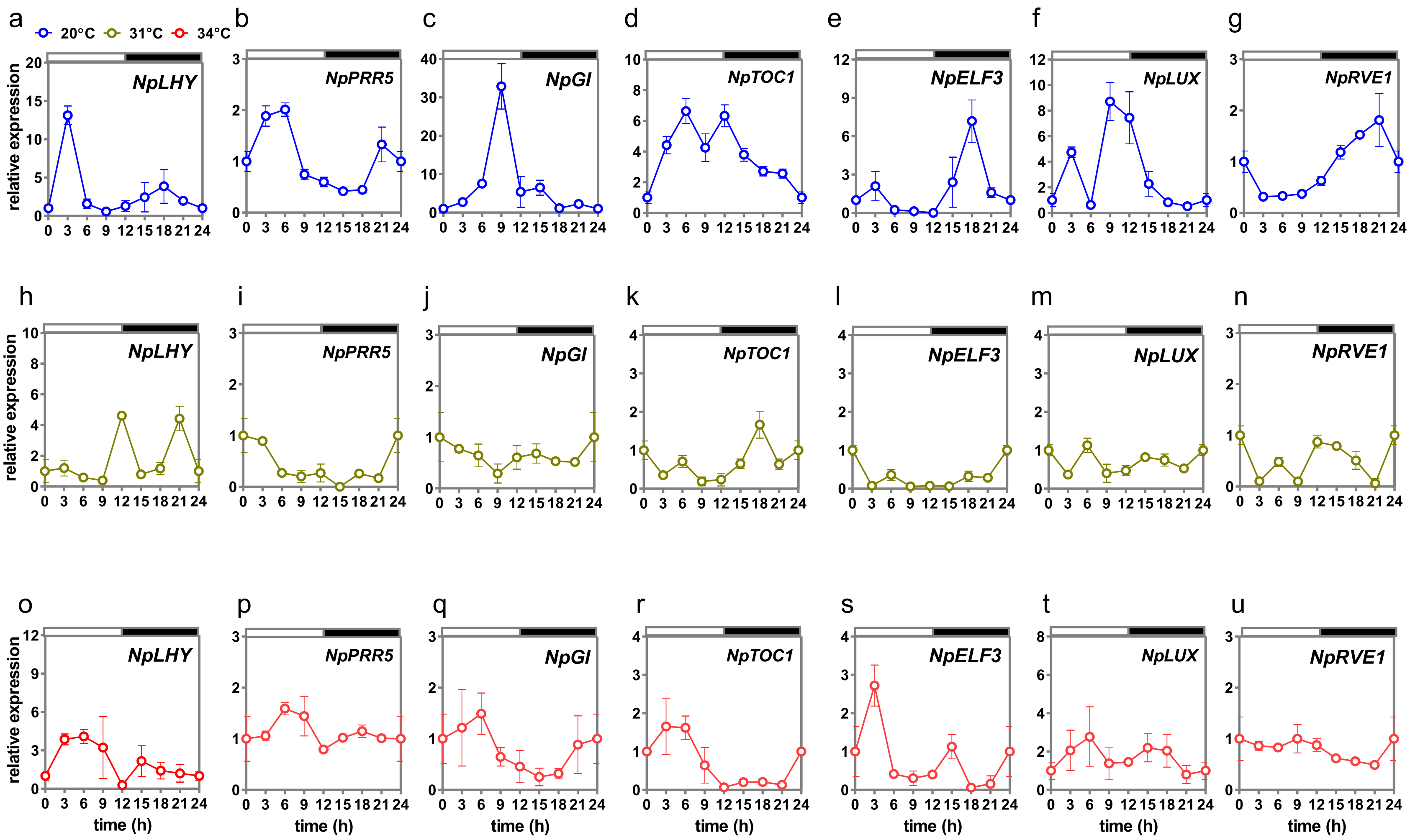


Fig. S7 | **Influence of temperature on the expression of clock genes under diurnal conditions in *N. pumilio***. Biological replicate of experiments shown in Fig. 5c-i. Seedlings were entrained with 12h light / 12h darkness photocycles at 20°C and then exposed to 20°C (blue, a-g), 31°C (brown, h-n) or 34°C (red, o-u). Samples were taken every 3h during the second day of each temperature treatment. Data represent mean and SD of three technical replicates. Gene expression levels at ZT0 were set to 1 and used as reference for all other time points. White and black rectangles represent day and night, respectively.

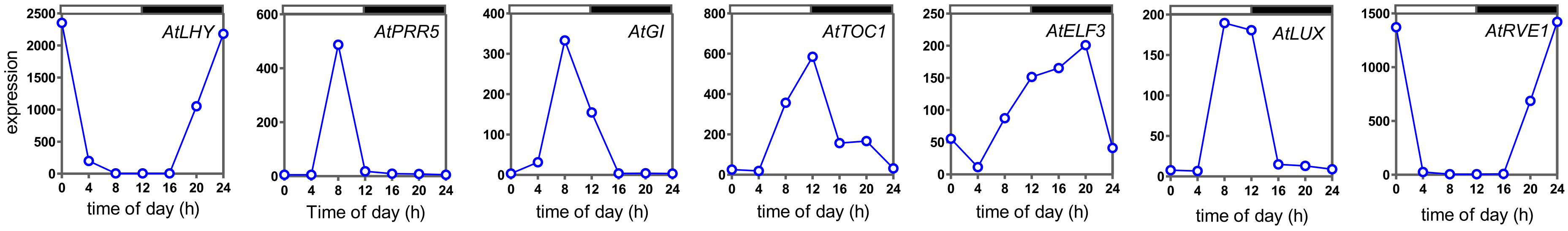


Fig. S8 | **Transcript levels of *Arabidopsis thaliana* clock genes AtLHY, AtPRR5, AtGI, AtTOC1, AtELF3, AtLUX and AtRVE1 under diurnal conditions**. Plants were entrained under photo-cycles (12h light / 12h darkness) at 20°C, and sampled every 4h under similar conditions. Data correspond to microarray experiments taken from DIURNAL (http://diurnal.mocklerlab.org/, LDHH_ST database). The complete experiment is described in Bläsing et al. 2005 (doi: 10.1105/tpc.105.035261).

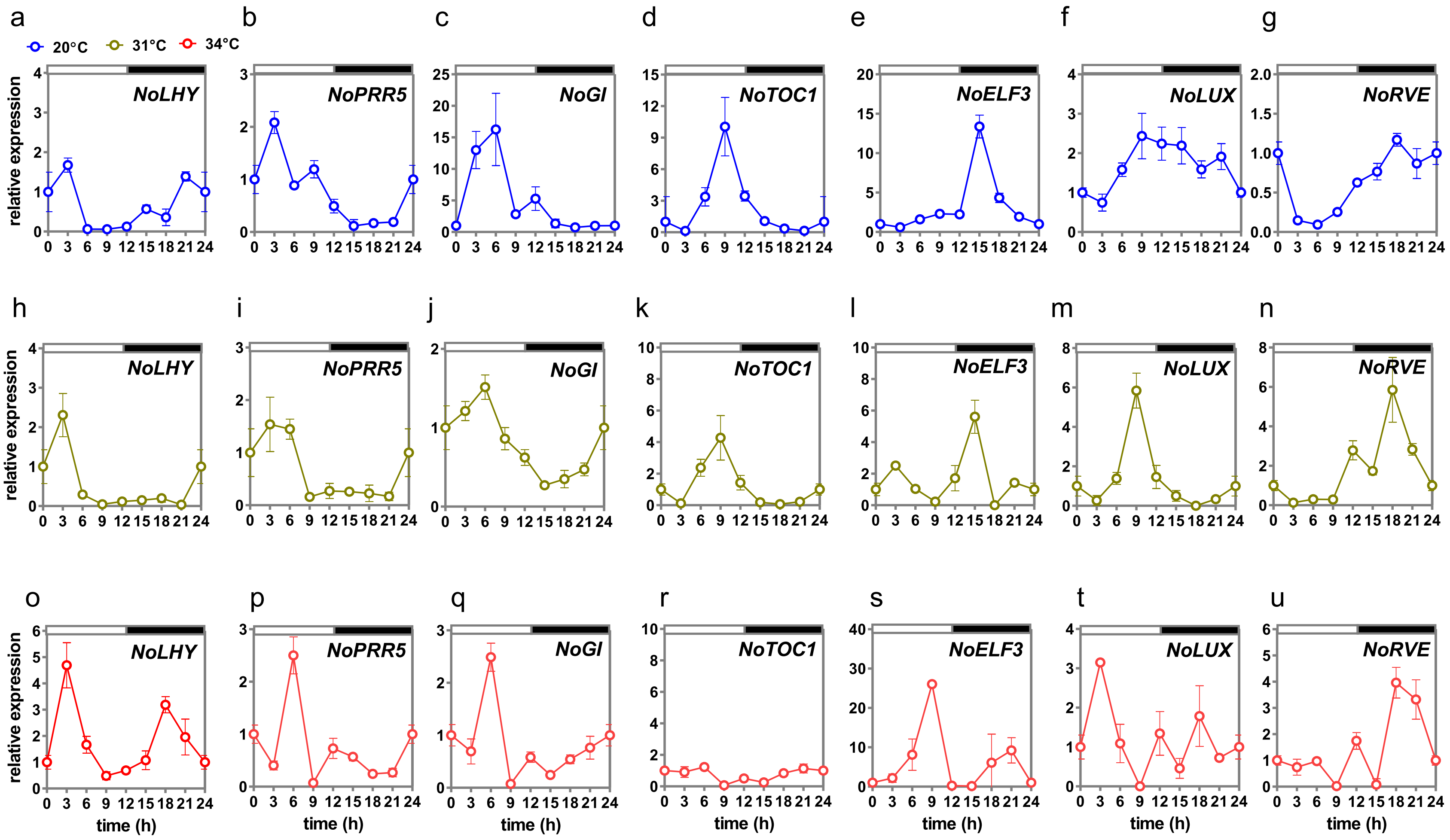


Fig. S9 | **Influence of temperature on the expression of clock genes under diurnal conditions in *N. obliqua***. Biological replicate of experiments shown in Fig. 5k-q. Seedlings were entrained with 12h light / 12h darkness photocycles at 20°C and then exposed to 20°C (blue, a-g), 31°C (brown, h-n) or 34°C (red, o-u). Samples were taken every 3h during the second day of each temperature treatment. Data represent mean and SD of three technical replicates. Gene expression levels at ZT0 were set to 1 and used as reference for all other time points. White and black rectangles represent day and night, respectively.

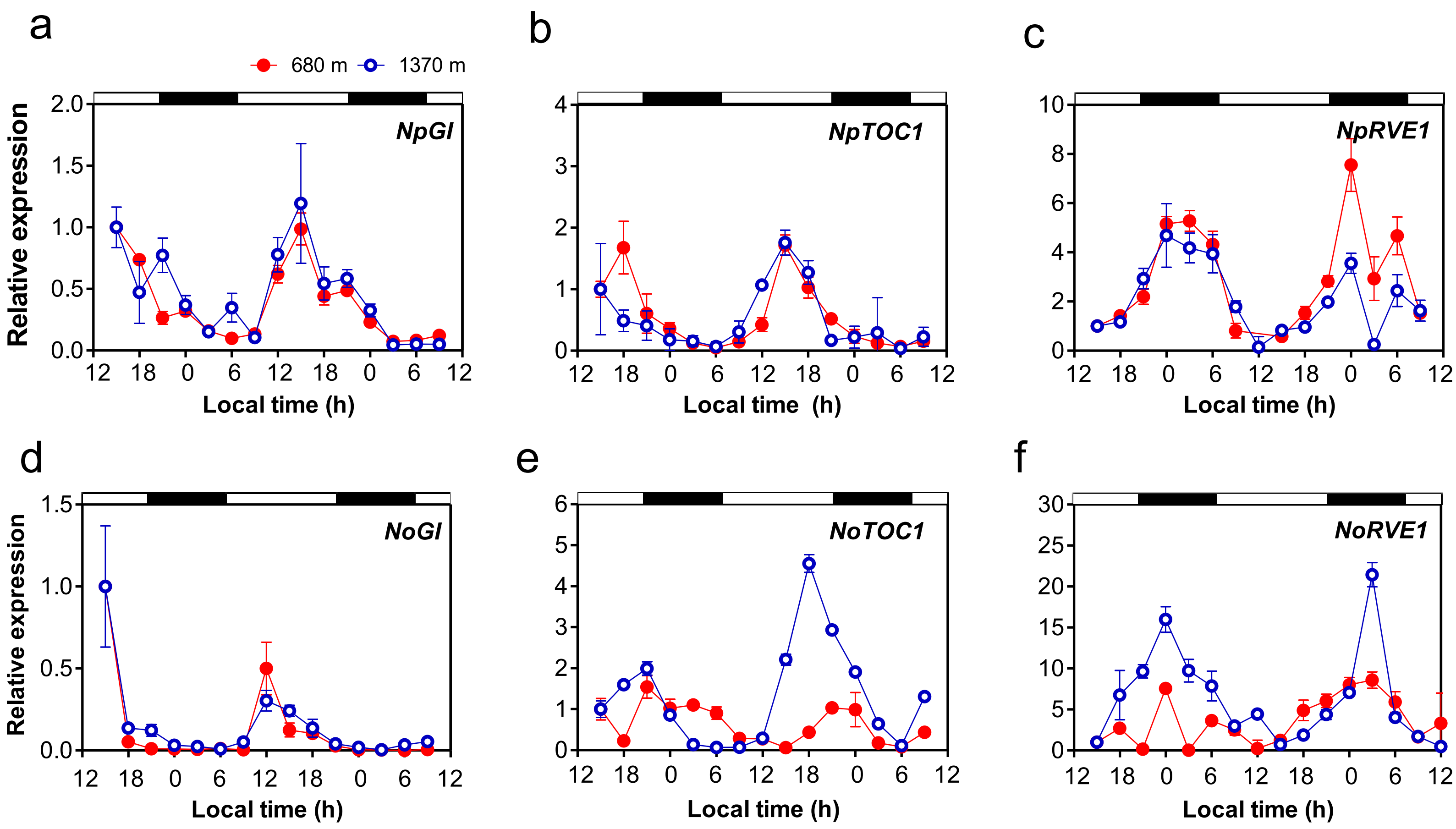


Fig. S10 | **Expression of core oscillator genes inside and outside the natural species' thermal ranges**. (a-c) Expression of *N. pumilio* clock genes *NpGI*, *NpTOC1*, *NpRVE1*. (d-f) Expression of *N. obliqua* clock genes *NoGI*, *NoTOC1*, *NoELF3* and *NoRVE1*. Data represent mean and SD of three technical replicates. Gene expression levels at time 0 were set to 1 and used as reference for all other time points. White and black rectangles represent day and night, respectively.

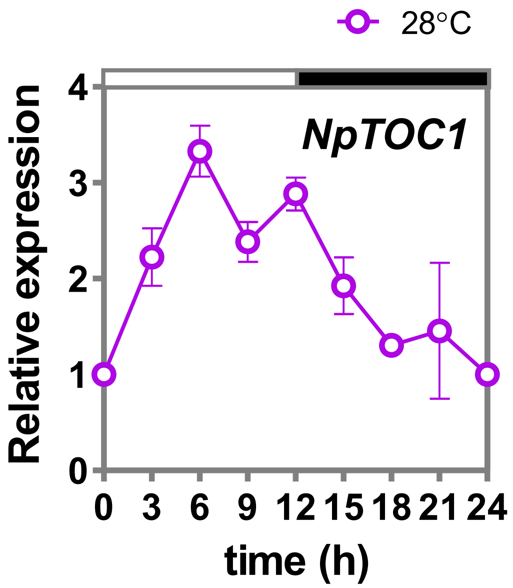


Fig. S11 | **Expression of *NpTOC1* at 28°C under diurnal conditions**. Seedlings were entrained with 12h light / 12h darkness photocycles at 20°C and then exposed to 28°C. Samples were taken every 3h during the second day of each temperature treatment. Data represent mean and SD of three technical replicates. Gene expression levels at ZT0 were set to 1 and used as reference for all other time points. White and black rectangles represent day and night, respectively

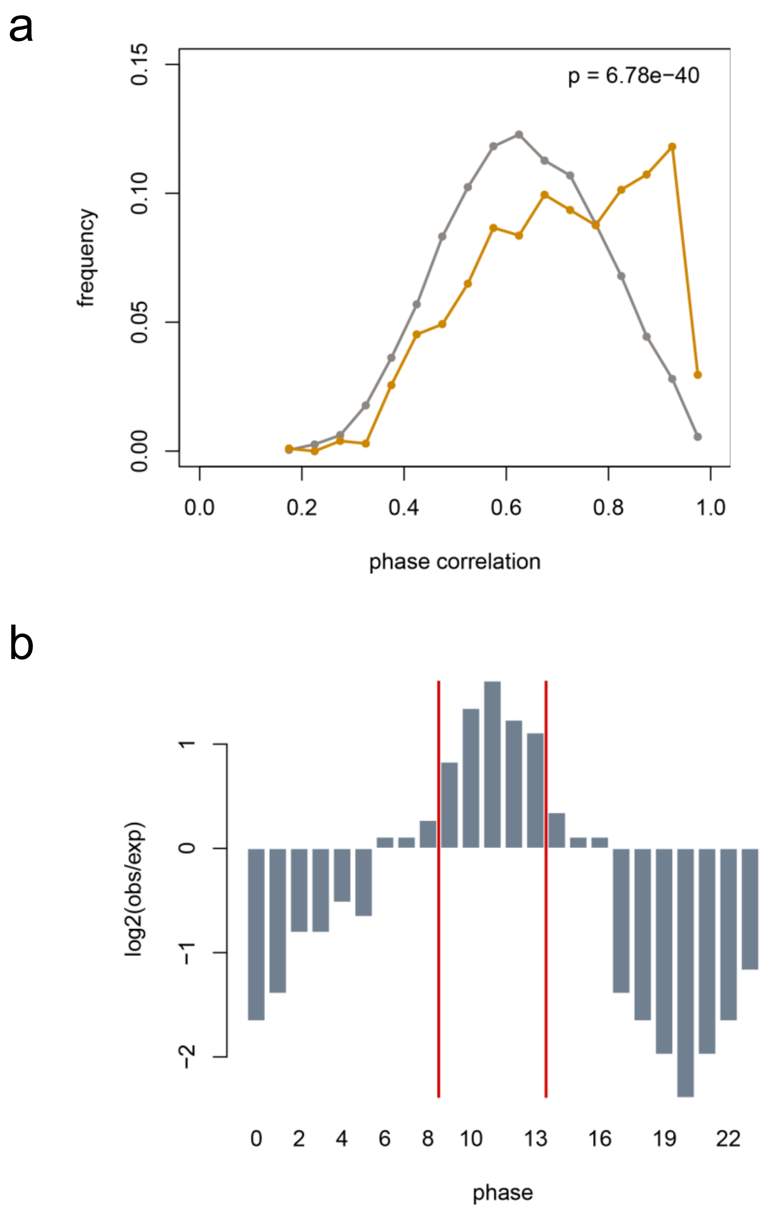


Fig. S12 | **Proof of concept of the method used for determining enrichment in oscillating genes and phase assignation used in Fig. 4**. Analysis of direct CCA1 targets genes described in Nagel et al. 2015 (www.pnas.org/cgi/doi/10.1073/pnas.1513609112). (a) Comparison between the expected (grey) and observed (orange) frequency distribution of genes with MBPMA values between 0 and 1. We used the MBPMA cut off of 0.8 as the threshold of cycling. (b) log2 fold change of observed vs expected ratio of the number of genes arranged by their time-of-day expression (phase). Phases of expression were assigned using the information of *Arabidopsis thaliana* genes that were strongly cycling (MBPMA cut-off of 0.8) of the database LL12_LDHH available in http://diurnal.mocklerlab.org/, and described in Harmer et al. 2000 (doi: 10.1126/science.290.5499.2110; see Methods).
